# Lamin B Receptor Upregulation in Metastatic Melanoma Causes Cholesterol-Mediated Nuclear Envelope Fragility

**DOI:** 10.1101/2023.12.21.572889

**Authors:** Michelle A. Baird, Robert S. Fischer, Cayla E. Jewett, Daniela Malide, Alexander X. Cartagena-Rivera, Clare M. Waterman

## Abstract

Metastatic cancer cells migrate through regions of tissue confinement, causing nuclear envelope (NE) rupture and heritable DNA damage. We discovered that cells from multiple cancers have increased NE fragility in confinement and transcriptional upregulation of nuclear genes compared to benign counterparts. A bioinformatic-driven siRNA screen revealed that lamin B receptor (LBR) upregulation correlates with melanoma progression and NE fragility. Increased LBR cholesterol synthase activity causes accumulation of cholesterol in the NE, which is necessary and sufficient for nuclear deformability and NE rupture in cells confined *in vitro* and is associated with NE rupture in invasive cells migrating out of tumor organoids and tumors *in vivo.* Thus, LBR upregulation causes excess NE cholesterol, driving nuclear fragility in confined migrating melanoma cells, establishing a direct role for nuclear membrane lipid composition in metastatic cancers.

---

Metastatic spread is responsible for the bulk of cancer patient deaths, nevertheless it remains a poorly understood process (*1–3*). During metastasis, tumor cells migrate through regions of cellular confinement, due to tightly packed local tissue architecture (*4*, *5*), remodeling of the extracellular matrix (*6*), and during trans-endothelial migration (*7*). While the cytoplasm is fluid and can easily deform through the 1-30µm pore size of such tissue regions (*8*), the nucleus is mechanically stiff, and its deformation during passage through small pores leads to rupture of the nuclear envelope (NE) (*9*, *10*). NE rupture exposes the genome to exonucleases in the cytoplasm (*9*, *11*, *12*), resulting in DNA damage and chromothripsis.

While abnormal nuclear size and morphology has long been used in cancer diagnosis (*13*, *14*), how this relates to the functional and mechanical properties of the NE or transcriptional changes that occur during oncogenesis is less understood. The NE is composed of two lipid bilayers bridged by nuclear pore complexes (NPC). The outer nuclear membrane (ONM) is contiguous with the endoplasmic reticulum (ER), while the inner nuclear membrane (INM) is enriched in a unique set of integral proteins that tether the lamin nucleoskeleton and heterochromatin to the nuclear periphery (*15*). Nuclear deformability and fragility are regulated by the interplay between the integral NE proteins of the INM (*16*), the heterochromatin and lamin nucleoskeleton (*17*, *18*), and via the cytoplasmic cytoskeleton which transmits forces to the NE (*19*). While lipid composition and distribution are known to alter membrane mechanical properties in many biological contexts (*20*), it remains unknown if the lipid composition of the ONM and INM are distinct, or if lipid composition affects NE functional or mechanical properties (*21*). Here, we sought to determine if changes in NE fragility and mechanics were an intrinsic property of malignant transformation, and if so, to address the mechanistic cause.

## NE genes are transcriptionally altered during cancer progression promoting NE fragility

We first examined whether expression levels of NE-associated genes were altered in cancer, and if this was related to nuclear mechanics and fragility. Utilizing the Human Protein Atlas (*22*), we generated an unbiased list of 249 NE genes and examined their expression levels in 6860 clinical samples representing 10 cancer subtypes in The Cancer Genome Atlas (TCGA) PanCancer repository (fig. S1A). We found altered expression levels of NE genes regardless of cancer type, supporting previous studies indicating dysregulation of nuclear-associated genes is an indicator of poor prognostic outcomes (*23*). To determine if dysregulation of NE genes correlates with confinement-induced NE rupture in cancer cells, we utilized high-resolution time-lapse live-cell imaging in conjunction with a polydimethylsiloxane (PDMS) cellular confinement device to mimic the degree of tissue confinement observed during metastasis *in vivo* (*24*). To assay NE permeability, we expressed mCherry fused to a nuclear localization signal (mCherry-NLS), which remains soluble in the nucleoplasm and leaks into the cytosol if the NE becomes permeable (*25*, *26*). To determine if NE rupture was extensive enough to expose chromatin to the cytoplasm, we expressed GFP-tagged cyclic GMP-AMP synthase (GFP-cGas), a component of the STING innate immune pathway, which is soluble in the cytosol and rapidly binds to dsDNA when chromatin breaches the NE (*27*), forming discrete fluorescent foci at the sites of exposure (Fig. 1A, B, S1B). We subjected a representative benign and malignant cell line (immortalized human melanocytes and the human metastatic melanoma cell line 1205Lu), to three different confinement heights (2.4µm, 3µm, and 5µm) to determine the degree of confinement required to induce NE damage (fig. S1C, D). We found that the nuclei of both the benign and malignant cells were insensitive to 5µm of confinement, while both cell lines showed nuclear fragility at 2.4µm of confinement. However, at the intermediate confinement height of 3µm there was a significant difference between the benign and malignant cell lines, with melanocytes maintaining NE integrity and 1205Lu exhibiting NE permeability and rupture.

**Figure 1.**
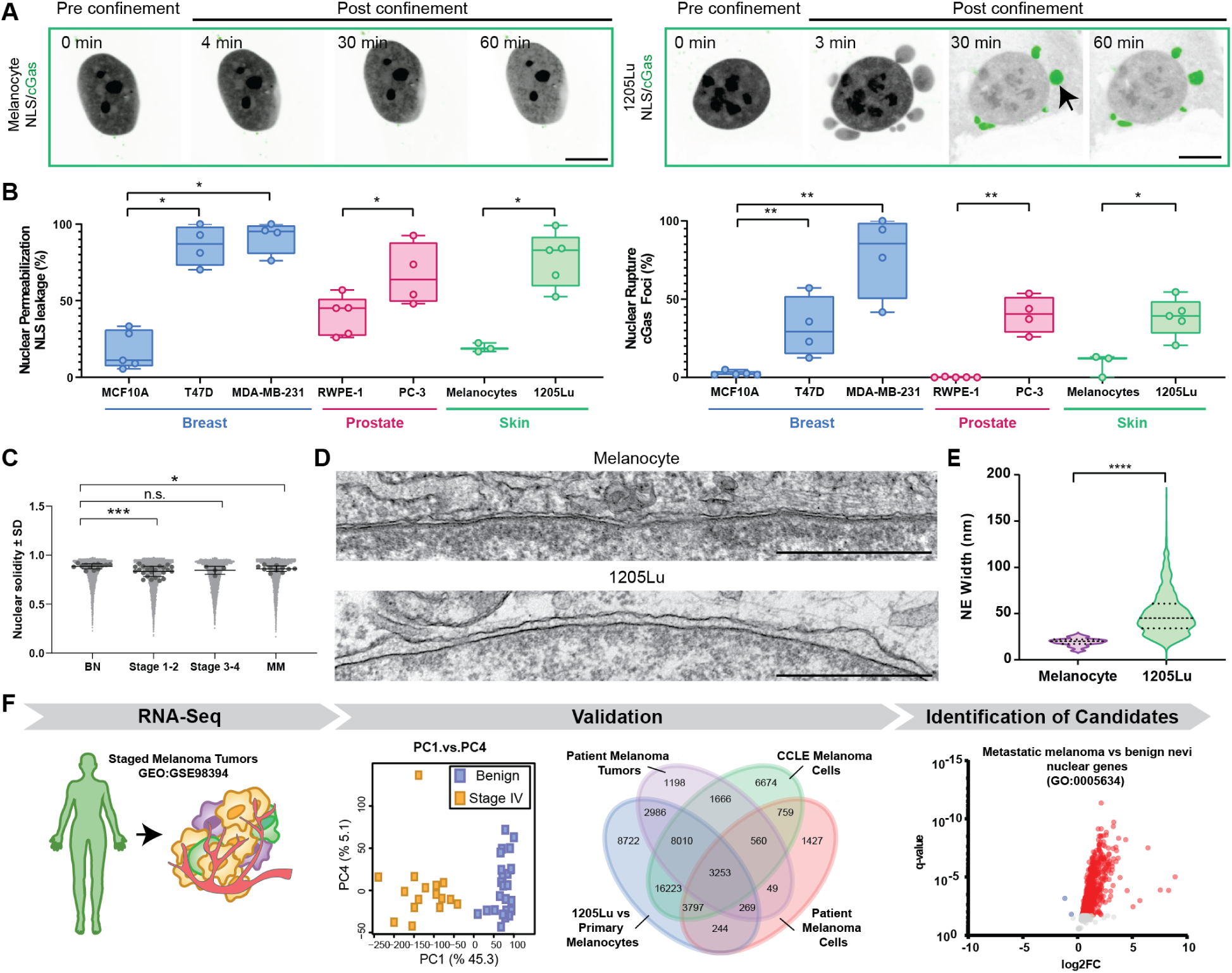
Transcriptional alterations in NE proteins correlate with melanoma disease progression and NE fragility. **(A)** Confocal image series of human melanocytes (left) or 1205Lu melanoma cells (right) transfected with mCherry-NLS (inverted grayscale) and GFP-cGas (green) before and after confinement to 3µm. Arrow highlights cGas focus. **(B)** Quantification of the fraction of confined cells exhibiting NE permeabilization (left) or NE rupture (right), for cell lines from benign and malignant breast (blue, MCF10A (benign, n=5 experiments, 332 cells), T47D (malignant, n=4 experiments, 187 cells), MDA-MB-321 (malignant, n=4 experiments, 192 cells), prostate (rose, RWPE-1 (benign, n=5 experiments, 124 cells), PC-3 (malignant, n=4 experiments, 153 cells) and skin, (green, melanocytes, benign, n= 3 experiments, 651 cells), 1205Lu (malignant, n= 5 experiments, 618 cells). Points= means of independent experiments. **(C)** Quantification of solidity (area/convex hull area) of nuclei in DAPI-stained tissue microarray samples from benign nevi (n= 3099 nuclei in 3 tissue samples), stage 1-2 (n= 3612 nuclei in 6 tissue samples), stage 3-4 (n= 1095 nuclei in 3 tissue samples), and malignant melanoma (n= 1055 nuclei in 3 tissue samples). Grey points=individual nuclei, black points= mean of each tissue sample **(D)** Representative EM micrographs of NEs in thin-sectioned human melanocyte (upper) or 1205Lu melanoma cells (lower) in tissue culture. Bar= 1µm. **(E)** Quantification of the gap between inner and outer NE from electron micrographs like those in (D), n= 400 measurements in 8 melanocytes, 866 measurements in 11 1205Lu cells. **(F)** Bioinformatic data analysis pipeline used to identify differences in transcript abundance in RNA-seq datasets (GEO:GSE98394) between patient biopsies from benign nevi and stage IV invasive tumors (left). Principal component analysis *(middle left)* confirmed distinct differences between nevus and stage IV. Genes whose transcript abundance differed (by Boneferroni post-hoc test) between nevus and stage IV datasets were cross-referenced against genes differentially expressed between human melanocytes and 1205Lu cells, as well as from the transcriptomes of 27 melanoma cell lines from the Broad cancer cell line encyclopedia and from FACS-sorted patient melanoma cells (GEO:GSE72056) (middle right, gene numbers in each subset shown). Genes that were present in all four datasets (3,253) were subjected to Gene ontology (GO) cellular component pathway analysis, and “nuclear associated” genes were subjected to *differential expression analysis* (volcano plot, right, red= significantly upregulated, blue= significantly downregulated, grey= non-significant). In (B, C), significance was tested with a Student’s T-test Mann-Whitney, (E), significance was tested with a Student’s T-test Kolmogorov-Smirnov, bars in B, E= median, in C= mean; boxes= 25^th^ to 75^th^ percentile, error bars in B= min and max, in C= S.D., in E= 25^th^ to 75^th^ percentile.

Importantly, 3D reconstructions of melanocytes and 1205Lu cells indicated no difference in unconfined nuclear height (fig. S1D). We then assayed cell lines representing benign and malignant breast and prostate tissues using a 3µm confinement height and our NE fragility biosensors (Movie S1). This showed that in cells from both tissue origins, the malignant cells had significantly more fragile NEs, as indicated by significantly greater presence of mCherry-NLS and GFP-cGas foci in the cytoplasm compared to their benign counterparts (Fig. 1A, B, S1B). Together, these results suggest that dysregulation of NE-associated genes and increased NE fragility under confinement are common intrinsic properties of malignant cells.

To determine if nuclear structure and nuclear-associated genes change as cancer progresses from benign to metastatic, we focused on melanoma, as its accessibility on the skin has facilitated availability of tissue samples and publicly available expression profile datasets from staged patient biopsies. Tissue microarrays from a cohort of 100 patients representing all clinical stages of melanoma were imaged and subjected to morphometric analysis of cell nuclei. This revealed significantly lower nuclear solidity in more advanced tumor samples, indicating nuclear lobularity increases with disease progression (Fig. 1C, S2A). To assess the NE in more detail, we performed transmission electron microscopy of immortalized human melanocytes and 1205Lu malignant melanoma cells (Fig. 1D). Analysis revealed that the spacing between the INM and ONM in benign cells was a consistent and uniform ∼45nm in width, while the melanoma cells displayed variable and inconsistent NE membrane spacing ranging from 4-177nm (Fig. 1D, E). Together these data indicate that alterations in nuclear morphology and NE structure are associated with malignant transformation of skin melanocytes.

We then utilized a bioinformatic approach to identify differential gene expression between benign and aggressive skin cancers. Employing RNA-seq transcriptomic datasets from biopsies of staged melanoma tumors (GEO:GSE98394) (*28*) (Fig. 1F), we focused on the two extremes of melanoma tumor progression, benign nevi and stage IV (metastatic melanoma). Given melanoma’s high mutational burden (*29*),we first applied principle component analyses to ensure the two datasets were distinct (Fig. 1F), followed by differential expression analysis (DEA) to identify a subset of genes exhibiting a significantly different number of normalized reads between nevi and stage IV samples (fig. S2B). To determine which transcriptional differences between the samples were consistent across many malignant melanomas, we cross-referenced this subset against genes differentially expressed between immortalized melanocytes and 1205Lu melanoma cells, as well as from the transcriptomes of 27 melanoma cell lines from the Broad cancer cell line encyclopedia (CCLE) (*30*), and from FACS-sorted patient melanoma cells (GEO:GSE72056) (*31*) (Fig. 1F). This resulted in 3,253 candidate genes that differed significantly in expression between benign and malignant melanoma (fig. S2B). We subjected this differentially expressed candidate list to cellular component gene ontology (GO) analysis, which demonstrated the second-most enriched category was nuclear-associated genes (fig. S2C), the majority of which were upregulated (Fig. 1F). Together, these results show that nuclear and NE abnormalities are associated with aggressive melanoma, and correlate with upregulation of nuclear-associated genes during disease progression.

## LBR upregulation is necessary and sufficient to promote NE fragility

We then sought to establish if transcriptional modulation of nuclear-associated genes affects NE fragility during cancer progression. As our bioinformatic analysis revealed that upregulation of nuclear-associated genes was more common than downregulation in advanced disease, and because upregulated proteins could potentially serve as biomarkers or drug targets, we selected a list of high priority upregulated gene targets representing a mix of known modulators of NE morphology and NE remodeling that also have strong correlation with cancer progression, (Fig. 2A). These included: Lamina-associated polypeptide 2b (Lap2β) and emerin, both of which bind to lamins and regulate gene expression and genome organization (*32*); Lamin B2, a nuclear intermediate filament protein of the lamin nucleoskeleton responsible for NE structural support and regulating genome organization (*33*); Nup93, a stable scaffold protein within the nuclear pore complex (*34*); CHMP2A, an interacting partner of endosomal sorting complex required for transport (ESCRT-III) protein that facilitates NE repair and re-sealing after mitosis (*35*); and lamin B receptor (LBR), an integral INM protein that binds chromatin and lamins and also mediates cholesterol biosynthesis (*36*). To determine if our priority upregulated NE genes affect NE fragility, we utilized siRNA to reduce their expression in 1205Lu cells. Cells were co-transfected with non-targeting (NT) or siRNAs targeting the gene of interest together with mCherry-NLS or GFP-cGas, subjected to confinement at 3µm, and the effects of knockdown (KD) on our markers of NE fragility were quantified, (Fig. 2B,C; S3A, B, Movie S2). This revealed that reduction of either Lap2β, CHMP2A, or laminB2 significantly decreased NE permeability (fig. S3B), while KD of either laminB2 or LBR nearly abolished NE rupture (Fig. 2C). Interestingly, changes induced by protein KD in NE permeability did not directly correlate with changes in NE rupture, indicating that NE permeabilization does not necessarily lead to catastrophic NE breakage, and *vice-versa*. Together, these results indicate that upregulation of components of the NE during melanoma progression increases NE fragility in confinement.

**Figure 2.**
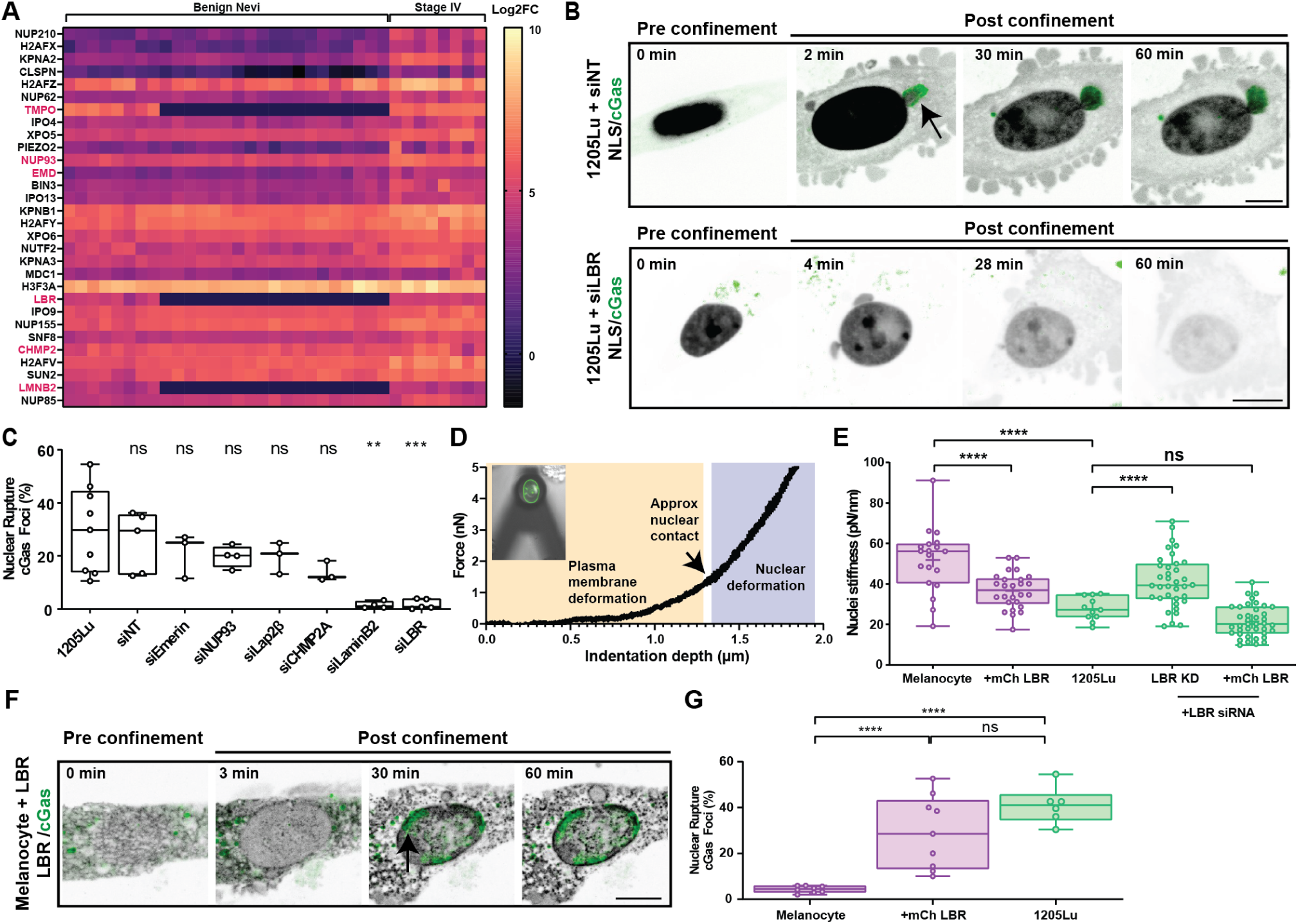
siRNA based screen identifies LBR as necessary and sufficient for promoting nuclear deformability and NE fragility under confinement. **(A)** Heat map of the top 30 nuclear-associated genes of interest based on transcript abundance from RNA-seq data from patient biopsies of benign nevi or stage IV invasive melanoma tumors, highlight indicates genes of interest chosen for siRNA-based screen. **(B)** Confocal image series of living 1205Lu melanoma cells transfected with mCherry-NLS (inverted grayscale) and GFP-cGas (green) together with either non-targeting siRNA, (siNT, upper) or siRNA targeting LBR (siLBR, lower) before and after confinement to 3µm. Arrow highlights cGas focus. **(C)** Quantification of the fraction of confined cells exhibiting NE rupture in 1205Lu cells with or without (1205Lu, n=9 experiments, 2473 cells) co-transfection with non-targeting siRNA (n=5 experiments, 421 cells) or siRNA targeting emerin (siEmerin, n=3 experiments, 261 cells), NUP93 (siNUP93, n=4 experiments, 521 cells), Lap2β (siLap2β, n=3 experiments, 195 cells), CHMP2A (siCHMP2A, n=3 experiments, 204 cells), LaminB2 (siLaminB2, n=4 experiments, 607 cells), or LBR (siLBR, n=5 experiments, 1627 cells). **(D)** Example AFM force curve indicating two distinct force regimes (membrane and cortex (peach) and nucleus (lavender) and transmitted light (greyscale) and fluorescence image (green) of a bead-affixed AFM cantilever resting on the nucleus of a living 1205Lu cell expressing GFP-Lap2β (inset). **(E)** Quantification of nuclear effective stiffness measurements of human melanocyte (purple) (n= 3 experiments, 19 cells) or 1205Lu cells (green) (n=3 experiments,11 cells) treated with LBR siRNA (LBR KD, n=4 experiments, 36 cells) or overexpressing mCherry-LBR (melanocytes + mCherry LBR n= 3 experiments, 25 cells; 1205Lu LBR KD +mCherry LBR n=5 experiments, 37 cells). **(F)** Confocal image series of a melanocyte co-transfected with mCherry-LBR (inverted greyscale) and GFP-cGas (green) before and after confinement to 3µm. Arrow highlights cGas focus. **(G)** Quantification of the fraction of confined cells exhibiting NE rupture (cGas foci) in 1205Lu cells (green, n= 6 experiments, 947 cells) or human melanocytes (purple) with (+mCherry LBR, n=8 experiments, 876 cells) or without expression of mCherry LBR (n=9 experiments, 1845 cells). In (B,F) bar = 10µm. In (C, E, G) points= means of individual experiments, In (C,G) significance was tested with a Student’s T-test Mann-Whitney, in (E) a student’s T-test using Welch’s correction, bars= means; boxes= 25^th^ to 75^th^ percentile, error bars= min and max.

To understand the mechanism by which NE gene upregulation promotes nuclear fragility, we chose to focus on LBR. LBR is an integral protein of the INM consisting of N-terminal TUDOR and RS domains that tether it to lamin B and heterochromatin (*37–39*) and a C-terminal 8-spanning transmembrane domain that contributes to cholesterol biosynthesis via its C-14 sterol reductase activity (*40*, *41*) (fig. S3C). Mutations in the C-terminal domain are associated with several human disorders, including Pelger-Huet anomaly which results in defects in neutrophil migration through small pores (*42*), and Greenberg skeletal dysplasia, a fatal prenatal disorder caused by abnormal sterol metabolism and developmental defects (*43*, *44*). Validation of our bioinformatics result by western blot confirmed that LBR was 30% more abundant in 1205Lu cells compared to immortalized melanocytes, and either pooled siRNA oligonucleotides or lentiviral-induced shRNAs targeting the 3’UTR of LBR (LBR-KD) reduced its level in 1205Lu cells to a similar level to that in immortalized melanocytes (fig. S3D). Importantly, re-expression of mCherry-LBR in wild type or LBR-KD cells showed that expressed LBR localized to clusters in the NE and in the ER (fig. S3E), as reported (*39*), and restored the confinement-induced cGas foci frequency to that of control cells (fig. S3F).

We then asked whether LBR contributes to nuclear mechanical properties by using simultaneous atomic force (AFM) and confocal microscopies to quantify nuclear stiffness and visualize nuclear shape. 1205Lu cells co-transfected with GFP-H2B to mark the nucleus and NT or LBR-KD siRNAs, either alone or together with mCherry-LBR, were indented 2.5µm with a bead-affixed AFM cantilever directly on top of their nuclei. This resulted in bi-modal force-displacement curves (Fig. 2D). The first mode was due to the cell membrane and cortex, while the second mode was sensitive to KD of laminA (*45*) and less sensitive to KD of laminB1 (*46*), and therefore represented the nuclear component (fig. S3G). Additionally, repeated indentations resulted in essentially identical force-displacement curves, indicating that our measurement was non-perturbative and reversible (fig. S3H). AFM mechanical analysis showed that compared to melanocytes, the nuclei of 1205Lu melanoma cells were much softer, and LBR-KD in 1205Lu caused nuclear stiffness to increase significantly to similar levels as melanocytes (Fig. 2E).

Importantly, re-expression of mCherry-LBR in LBR-KD cells softened nuclei to levels similar to that in wild type 1205Lu (Fig. 2E). We next asked if increasing LBR alone was sufficient to alter nuclear mechanics and susceptibility to rupture in non-malignant cells. Overexpression of mCherry-LBR in immortalized melanocytes significantly decreased the stiffness of nuclei to a level comparable to that in 1205Lu (Fig. 2E). Furthermore, our confinement assay and nuclear fragility biosensors in immortalized melanocytes showed that overexpression of LBR was sufficient to increase the frequency of NE rupture by more than 3.5-fold, raising it to the level observed in 1205Lu cells (Fig. 2F, G, Movie S3). This shows that high LBR expression in metastatic melanoma is necessary and sufficient not only to render the NE susceptible to damage under confinement, but also to modulate nuclear deformability, generating a softer, more fragile nucleus.

We then sought to determine the cellular mechanism by which excess LBR causes chromatin to breach the lamin nucleoskeleton, the INM, and the ONM during confinement-mediated NE rupture. We first analyzed the effect of LBR-KD on the organization of proteins involved in NE function or nuclear mechanics. We found that LBR-KD in 1205Lu cells had little effect on either gross nuclear architecture or distribution of proteins associated with the LINC complex (Sun2), nuclear pores (mab414), the lamin nucleoskeleton (lamins A/C, B1 and B2), or condensed heterochromatin (H3K9me2, H3K9me3, H4K20me3), although it did cause minor reduction in Lap2β at the NE (fig. S4A, B). We then examined the fate of the lamin nucleoskeleton during confinement-induced NE rupture by immunostaining laminA in 1205Lu cells co-expressing mCherry-NLS and GFP-cGas. In confined cells displaying mCherry-NLS in the cytoplasm and perinuclear cGas foci indicative of nuclear rupture, we found that laminA staining was absent from the site where DNA protruded from the nucleus into the cytoplasm and cGas foci formed, indicating breach of the nucleoskeleton (fig. S4C). We next probed the fate of the ONM and INM during NE rupture by expressing our nuclear permeability biosensor together with GFP-Lap2β to mark the INM, SiR staining of DNA, and an ER vital dye to mark the ONM. Analysis of superresolution images of 1205Lu cells during confinement revealed large NLS-containing containing blebs that formed at the surface of the nucleus and which were enclosed in ER-positive membrane that was contiguous with the NE, indicating it was an extension of the ONM (Fig. 3A, Movie S1). These ONM-encased blebs grew and filled with mCherry-NLS and DNA over time until the ONM ruptured, releasing mCherry-NLS into the cytosol (Movies S1, 2).

**Figure 3.**
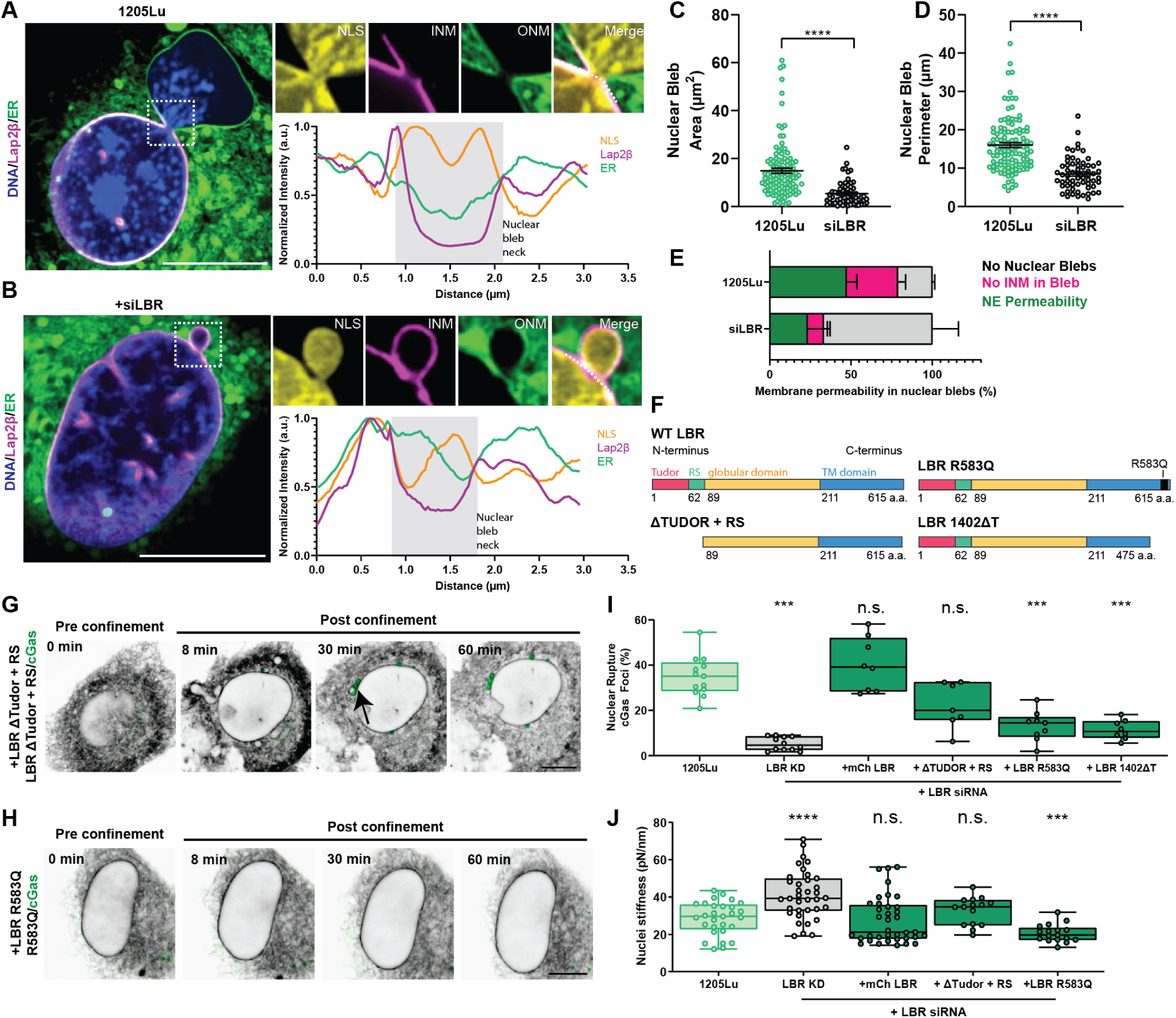
LBR sterol reductase activity promotes INM fragility to mediate NE rupture under confinement. **(A,B)** Super-resolution confocal images taken during confinement to 3µm of living 1205Lu melanoma cells transfected with mCherry-NLS (yellow) and GFP-Lap2β (magenta) and stained with blue-white ER-Tracker (green) and SiR DNA (blue); with (B) or without (A) transfection with siRNAs targeting LBR (+siLBR). Zooms of boxed regions (above, right), intensity linescans of dotted lines (below, right). **(C, D)** Quantification nuclear bleb area (C) and perimeter (D) from images of cells treated as in (A, B) (1205Lu: green, n=3 experiments, 116 cells; siLBR: gray, n=3 experiments, 137 cells) **(E)** Quantification of the fraction of cells exhibiting no nuclear blebbing (gray), no INM within nuclear blebs (pink) or NE permeability (green) from time-lapse movies of cells treated as in (A, B) (1205Lu: n=4 experiments, 4 cells; siLBR: n=4 experiments, 4 cells). **(F)** Schematic diagram of LBR functional domains and mutants. **(G, H)** Confocal image series before and after confinement to 3µm of 1205Lu cells co-transfected with GFP-cGas (green) and siRNA targeting the 3’UTR of LBR together with either mCherry-LBRDTudor+RS (G, inverted grayscale) or mCherry LBR 583Q (H, inverted grayscale). Arrow highlights cGas focus. **(I)** Quantification of the fraction of confined cells exhibiting NE rupture in 1205Lu cells expressing GFP-cGas alone (light green, 1205Lu, n= 6 experiments, 746 cells) or together with LBR siRNA (gray, LBR KD, n=4 experiments, 3283 cells) and either mCherry LBR (dark green, +mCherry LBR n=3 experiments, 1342 cells), mCherry-LBRDTudor+RS (dark green, n=3experiments, 610 cells), mCherry-LBR 583Q (dark green, n=3 experiments, 1142 cells) or m-Cherry-LBR-Y468T (dark green, n=3 experiments, 228 cells). **(J)** Quantification of nuclear stiffness in 1205Lu cells (light green, n=5 experiments, 30 cells) expressing LBR siRNA (gray, LBR KD, n=4 experiments, 36 cells) and either mCherry LBR (dark green, n=5 experiments, 37 cells), mCherry-LBRDTudor+RS (dark green, n=3 experiments, 15 cells), or mCherry-LBR 583Q (dark green, n=3 experiments, 17 cells). Data points represent individual cells (C, D, J) or independent experiments each consisting of pooled cell data from a single imaging dish (I). In (A, B, G, H) bar = 10µm. In (C, D, I) significance was tested with a Student’s T-test Mann-Whitney, in (J) a student T-test using Welch’s correction. Bars=means, in (C,D,I,J) error bars=min and max, in (E) error bars=SEM

Comparison of the INM marker GFP-Lap2β and the ER ONM marker showed that a small discontinuity in the INM formed locally concomitant with the establishment of the ONM bleb (Fig. 3A, Movie S4), which remained largely devoid of INM marker as it expanded, filled with mCherry-NLS and DNA, and subsequently burst. In contrast, NE membrane blebs in confined LBR-KD cells were less frequent, smaller, and were generally encased by both the INM and ONM (Fig. 3B-E, Movie S4). Together these results indicate that upregulation of LBR in melanoma promotes fragility of the INM, which upon confinement, ruptures and allows nuclear pressure to drive formation of large ONM blebs containing nucleoplasm and chromatin, which then burst, resulting in nuclear DNA exposure to the cytoplasm.

## LBR sterol reductase activity promotes INM fragility

Our next goal was to uncover the molecular mechanism mediating LBR’s promotion of nuclear softening and NE fragility. We generated mCherry-tagged LBR variants (Fig. 3F, S4D), including a truncation mutant that lacked the N-terminal lamin and chromatin binding domains (mCherry-LBR-ΔTudor+RS) and two human disease-related point mutants (mCherry-LBR-R583Q and mCherry-LBR-1402ΔT), both of which result in abolishment of the sterol reductase activity and which are associated with Greenberg’s skeletal dysplasia (*40*, *47*, *48*). Analysis of NE rupture in confinement and nuclear deformability by AFM of LBR-KD cells expressing mCherry-LBR-ΔTudor+RS revealed that this mutant rescued the effects of LBR-KD on nuclear deformability and cGas foci formation, similar to LBR-KD cells expressing mCherry-LBR (Fig. 3G, I, J, Movie S8). Thus, chromatin tethering by LBR is dispensable for effects on nuclear mechanics and NE fragility. Similarly, AFM analysis of LBR-KD cells expressing mCherry-LBR-R583Q showed that this mutant rescued the decrease in nuclear deformability caused by LBR depletion, inducing nuclear softening to a level even greater than that of either 1205Lu cells or LBR-KD cells rescued with mCherry-LBR (Fig. 3H, J). Surprisingly, neither mCherry-LBR-R583Q nor mCherry-LBR-1402ΔT could rescue the decrease in NE rupture under confinement caused by knockdown of LBR (Fig. 3H, I, Movie S5). This demonstrates that the C-14 sterol reductase activity of LBR specifically promotes NE fragility in melanoma cells under confinement, and further shows that nuclear deformability and NE fragility are governed by mechanistically distinct functions of LBR.

## LBR drives NE fragility through NE cholesterol accumulation

To determine how LBR’s role in cholesterol synthesis promotes NE fragility under confinement, we examined the contribution of LBR to total cellular cholesterol and asked whether LBR contributes cholesterol specifically to the NE. Fluorometric analysis of 1205Lu or LBR-KD cells alone or re-expressing either mCherry-LBR or mCherry-LBR-R583Q cultured in lipid-free media showed that LBR-KD significantly reduced cholesterol, and rescue of LBR-KD with wild type LBR restored cholesterol almost to control level, while rescue with the LBR sterol reductase mutant did not (Fig. 4A). To find out if LBR specifically contributes to cholesterol enrichment in the NE or within nuclear blebs, we performed super-resolution microscopy of cells bearing manipulations of their LBR level or activity that had been stained with bodipy-cholesterol, SiR-DNA and ER vital dye to localize cholesterol, chromatin, and the ONM, respectively (Fig. 4B, D, F, H, S5A). This showed in 1205Lu cells that cholesterol was present in the plasma membrane, membranous organelles, lipid droplets, as well as the NE and ER (Fig. 4B, S5A). In contrast, LBR-KD cells exhibited a strongly reduced intensity of bodipy cholesterol throughout these regions compared to 1205Lu cells (Fig. 4B). Analysis of the NE revealed high and homogeneous cholesterol distribution in the NE of 1205Lu cells, resulting in a low variance in fluorescence intensity along the NE, while LBR-KD cells exhibited cholesterol clusters interspersed by areas lacking cholesterol, resulting in a significantly higher intensity variance along their NE (Fig. 4C,D). Pairwise comparison of regional bodipy cholesterol fluorescence within individual cells showed in 1205Lu cells that cholesterol was significantly higher in the NE compared the ER and was highly concentrated at the base of nuclear blebs compared to the non-bleb NE, and these local differences were abrogated by LBR-KD (Fig. 4F-I). Together, these results indicate that upregulation of LBR in melanoma cells increases cellular cholesterol, which concentrates in the NE and accumulates at sites of NE blebbing and rupture.

**Figure 4.**
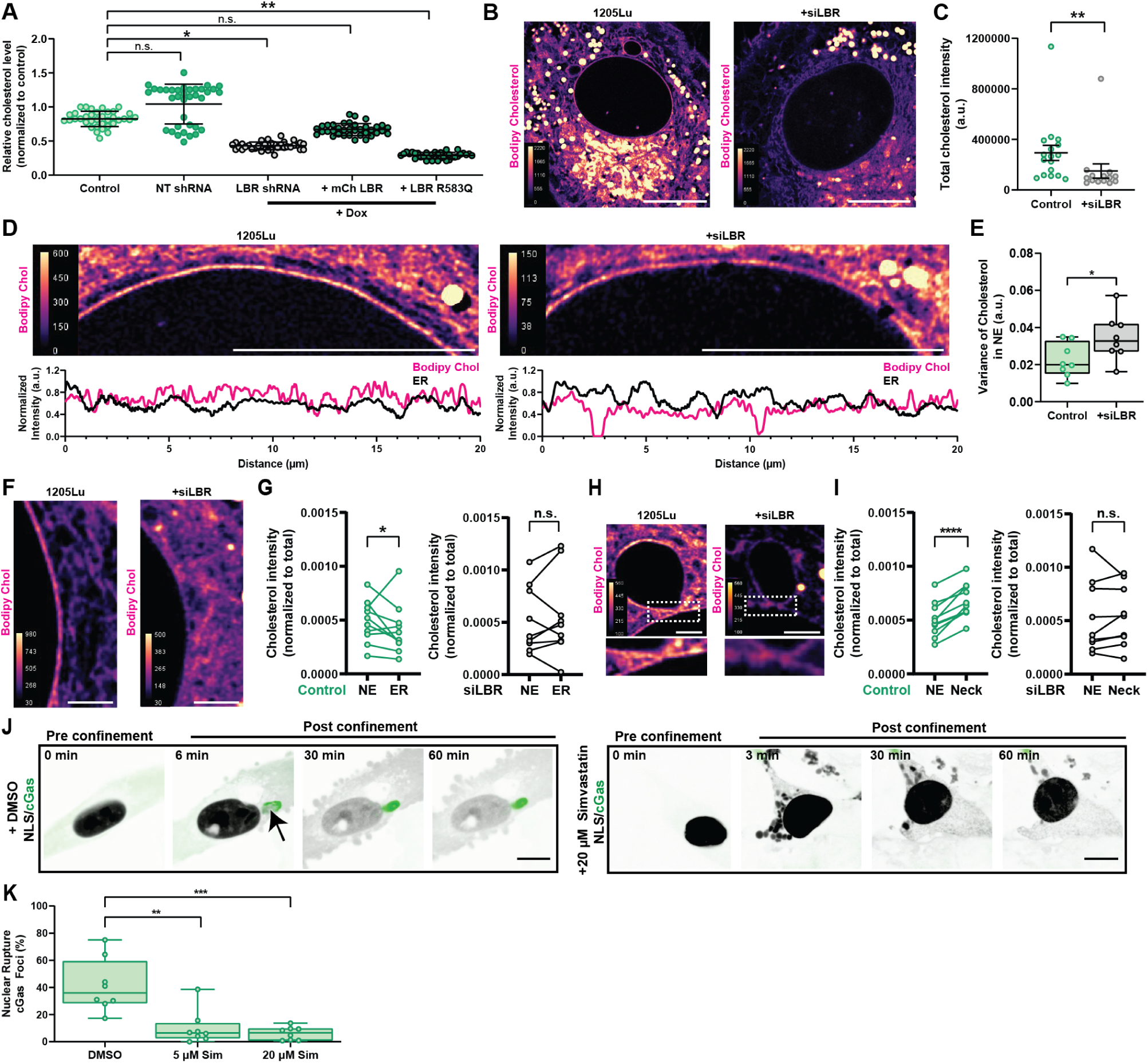
LBR-mediated cholesterol accumulation in the NE drives nuclear rupture in confinement. **(A)** Spectrophotometric analysis of cholesterol level in of 1205Lu (Control) with induced (+ Dox) expression of either non-targeting shRNA (NT shRNA) or shRNA targeting the 3’UTR of LBR (LBR shRNA) alone or together with either mCherry fused to wild type (+mCh-LBR) or mutant LBR (+ LBR R583Q). n= 3 experiments of 12 replicates each. **(B, D, F, H)** Intensity color-encoded super-resolution confocal images taken during confinement to 3µm of 1205Lu melanoma cells with (+siLBR, right) or without (1205Lu, left) transfection with siRNAs targeting LBR and stained with bodipy cholesterol. Note that color scale bars are the same between conditions in (B, H) to highlight difference in total staining, while color scale bars are different between conditions in (D, F) to highlight variation in staining along the NE (D, linescan below) and difference in relative staining of the ER and NE (F). **(C)** Quantification of total bodipy cholesterol staining per cell from images like those in (B). n= 17 cells (control) 14 cells (LBR siRNA) in 3 experiments. **(E)** Quantification of variance in intensity measured along 2µm long line-scans of the NE taken from images of bodipy cholesterol-stained cells like those in (D). n=3 experiments in 8 cells each condition. **(G, I)** Quantification of normalized (to total) bodipy cholesterol intensity measured for equal areas of NE and ER (G, n= 3 experiments, 10 cells for control, 9 cells for LBR siRNA) or of NE and at the base of nuclear blebs (Neck, like boxed regions in (H)) (I, n= 3 experiments, 10 cells for control, 9 cells for LBR siRNA), taken from images of bodipy cholesterol-stained 1205Lu cells like those in (F, H), respectively. Paired points represent measurements taken within the same cell. **(J)** Confocal image series of 1205Lu cells treated with vehicle (+DMSO) or 20µM simvastatin and expressing mCherry-NLS (inverted grayscale) and GFP-cGas (green), before and after confinement to 3µm. Arrow highlights cGas focus. **(K)** Quantification of the fraction of confined cells exhibiting NE rupture in 1205Lu cells treated with vehicle (+DMSO, n=3 experiments, 443 cells) or 5 (n=3 experiments, 1182 cells) or 20µM (n=3 experiments, 559 cells) simvastatin and expressing GFP-cGas. Data points represent individual cells (C, E,G,I), or independent experiments each consisting of pooled cell data from a single imaging dish (K), or means of individual experiments (A). In (B,D) bar = 10µm, in (F, H) bar= 2mm. In (A) significance was tested with ANOVA, in (A), Student’s T-test with Kolmogorov-Smirnov correction (C), T-test with Welch’s correction (E), paired T-tests (G,I), and T-test Mann-Whitney correction (K). Bars in (A, C, E, K)= mean; in (A, C) error bars=SD. In (E,K) error bars= min and max, in (K, E) boxes= 25^th^ to 75^th^ percentile.

We next asked whether reduction of cellular cholesterol was sufficient to prevent NE rupture in confined melanoma cells. We utilized the statin, simvastatin, which inhibits HMG CoA reductase, an enzyme upstream of C14 sterol reductase in the cholesterol biosynthetic pathway (*49*). Simvistatin treatment reduced cellular cholesterol in 1205Lu cells (fig. S5B). Pretreatment of 1205Lu cells expressing our NE fragility biosensors with simvastatin followed by confinement revealed that while treatment induced marked changes in nuclear morphology under confinement, with nuclei appearing much more deformable and fluid-like, treated cells exhibited a significant decrease of GFP-cGas foci compared to untreated cells (Fig. 4J, K, Movie S6). This indicates that cholesterol reduction is sufficient to render melanoma cells resistant to NE rupture in confinement. Together, these results support a mechanism whereby LBR-mediated increase in cholesterol in melanoma cells causes cholesterol accumulation in the NE that renders the INM fragile and susceptible to rupture in confinement.

## LBR promotes NE rupture during metastasis *in vivo*

Finally, to test the role of LBR in tumor progression and NE fragility in physiologically relevant models, we generated 1205Lu cell lines stably expressing GFP-cGas along with inducible LBR- or NT-shRNAs (fig. S6A), and generated tumor organoids *in vitro* and intradermal tumors in mice. Organoids were embedded in collagen matrices and allowed to proliferate and invade into the surrounding ECM for 1 week prior to 3D confocal imaging (Fig. 5A, S6C). Analysis showed that depletion of LBR had no significant effect on organoid volume (fig. S6B), but significantly decreased both the number of cells that detached from the tumor and were present in the ECM (Fig. 5B) and the distance they travelled from the organoid centroid compared to WT 1205Lu and NT-shRNA (Fig. 5C). Although the high cell density in organoid cores made it impossible to accurately measure NE rupture, we were able to quantify perinuclear GFP-cGas foci in a 250mm-thick 3D shell around the organoid edge as well as in cells outside this zone that had invaded the ECM. This revealed a low base level of NE rupture in the organoid edge, present in all conditions (Fig. 5D). However, in the invasive periphery, while WT or NT organoids displayed many invasive cells bearing GFP-cGas foci, LBR-KD organoids showed significantly less cells with this indication of NE rupture (Fig. 5A,D, S6C,D, Movie S7,8), despite nuclei appearing deformed and confined and similar collagen meshwork density and architecture around organoids in all conditions (fig. S6E,F). Analysis of the minimal diameter of nuclei as a proxy for nuclear deformation showed that those nuclei with associated GFP-cGas foci were significantly skinnier than those without, independent of condition (Fig. 5E), suggesting that rupture was specific to nuclei subjected to confinement by the ECM or other cells. To determine whether LBR promotes NE fragility and rupture in skin tumors *in vivo,* we injected 1205Lu cells expressing LBR- or NT-shRNA together with the GFP-cGas biosensor intradermally into mice and allowed tumors to grow and invade for two months (Fig. 5F, S6G). Harvesting tumors for high resolution 2-photon imaging *ex vivo* revealed substantial ECM rearrangement and melanoma cell migration in all experimental conditions, as indicated by the presence of thick linear bands of collagen peripheral to the tumor edge and cells appearing to use both single and collective migratory modes to exit the primary tumor (Fig. 5F, S6E). Analysis showed that depletion of LBR reduced tumor volume compared to controls, however didn’t rise to statistical significance (fig. S6I). Although we were unable to accurately quantify the absolute number of perinuclear cGas foci because of resolution limitations and background fluorescence deep in tissue, we did note that nuclei-associated GFP-cGas foci occurred predominantly at the tumor periphery, specifically in cells metastasizing through dense collagen, and nuclear rupture appeared more frequent in control tumors compared to LBR-KD (Fig. 5F, S6G, H, Movie S9,10). Quantification of minimal nuclear radius in tumors *in vivo* revealed that nuclei with cGas foci were significantly skinnier than those without foci, independent of condition (Fig. 5G), implying that cells positive for NE rupture were experiencing local confinement. Together, these data demonstrate that excess LBR in aggressive melanoma promotes tumor growth, metastasis and NE fragility specifically during conditions of invasive, confined migration.

**Figure 5.**
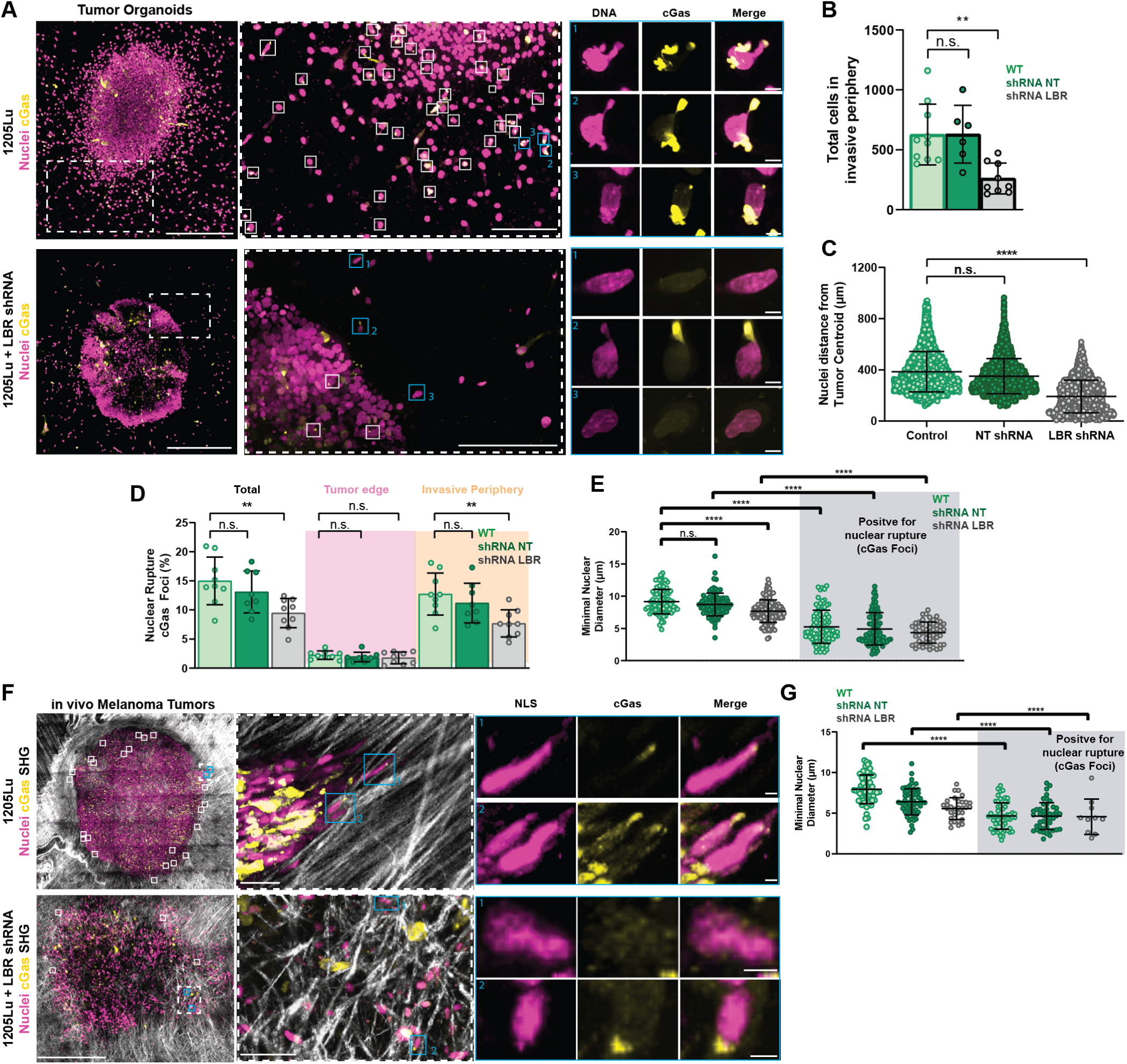
LBR promotes nuclear rupture in cells migrating out of tumor organoids and epidermal tumors *in vivo*. **(A)** 3D projections of z-series of confocal images of fixed tumor organoids grown from 1205Lu cells stably expressing GFP-cGAS (yellow) and mScarlet-NLS (magenta) with (+LBR shRNA) or without (1205Lu) stable expression of shRNA targeting LBR for 21 days in a collagen ECM. Dotted boxed regions from left are zoomed (left center), white boxes indicate cells positive for cGas foci, blue boxed regions from center are zoomed (right center). **(B-E)** Quantification of total number of cells in the invasive periphery (B) distance of nuclei from the organoid center (C), percent of nuclei exhibiting rupture in the total organoid and at the tumor edge (dusky rose) and invasive periphery (gold) (D), nuclear diameter with (grey, right) and without (white, left) cGas foci for organoids grown from 1205Lu cells stably expressing GFP-cGAS and mScarlet-NLS with or without (WT, green) stable expression of non-targeting shRNA (shRNA NT) or shRNA targeting LBR (+LBRshRNA) for 21 days in a collagen ECM. **(F)** 3D projections of z-series of two-photon images of live tumors imaged *ex vivo* grown from 1205Lu cells stably expressing GFP-cGAS (green) and mScarlet-NLS (magenta) with (+LBRshRNA) or without (1205Lu) stable expression of shRNA targeting LBR and labeled with hoechst. Solid boxed regions (left) represent cells positive for cGas foci. Dotted boxed region from left are zoomed (left center) and blue boxes indicate zoomed nuclei (right). **(G)** Quantification of tumor volume and **(H)** nuclear diameter with (grey, right) and without (white, left) cGas foci. Data points represent individual cells (C,E, H,) or independent experiments each consisting of pooled cell data (B,D, G,). In (A) bar = 500µm (far left), 150µm (middle) and 5µm (far right). In (F) bar = 500µm (far left), 50µm (middle) and 5µm (far right).In (E,H) significance was tested with ANOVA (,D,) significance was tested with a Student’s T-test Mann-Whitney, in (C) a student T-test using Welch’s correction. Bars= means with SD, in (B,C,D,E,G,H), (D) error bars= min and max boxes= 25^th^ to 75^th^ percentile.

We show here that upregulation of nuclear membrane proteins occurs as cancer progresses from benign to metastatic, promoting nuclear deformability and NE fragility in the context of 3D confined cancer cell migration, as seen during cancer dissemination. Focusing our efforts on a mechanistic understanding of LBR, we discovered that upregulation of LBR during melanoma progression leads to concentration of cholesterol in the NE, promoting nuclear deformation and INM fragility through local modulation of NE lipid composition. We suggest that upregulation of LBR during melanoma progression could be playing dual roles. First, by increasing nuclear deformability to ease the impediment a large, stiff nucleus poses to cancer cell migration through confining tissues, however at the cost of increasing NE fragility and thereby facilitating DNA damage and genomic instability. Secondly, LBR’s role in cholesterol synthesis could serve as a metabolic enhancer, increasing tumor cells’ ability to proliferate (*50*) and cope with nutrient-deprived conditions (*40*). We propose a model (fig. S7), where LBR-mediated accumulation of cholesterol in the NE generates cholesterol islands that may cause regional defects in membrane flow, where if exposed to mechanical stress such as migration through confining spaces, the INM is unable to flow to expand, allowing buildup of pressure and eventual failure, resulting in localized loss of NE integrity, exposing chromatin to the exonuclease-rich cytoplasm, thus generating DNA damage and genetic instability. This is particularly intriguing in the context of cancer, as high cholesterol has been associated with tumor development (*51*) and the activity of tumor infiltrating T cells in melanoma (*52*), and meta-analysis of individuals has shown that long term statin use is associated with decreased cancer progression and severity in many cancer subtypes (*53*), including melanoma (*54*, *55*). Together, these data establish a direct connection between upregulation of NE genes capable of modulating nuclear membrane lipid composition in the promotion of NE fragility in metastatic cancers, and provide a novel category of biomarkers of melanoma progression and potential therapeutic drug targets.

## Acknowledgments

We thank the Waterman Lab and Dr. Glenn Merlino for many vibrant discussions on this work, as well as the Physiology Course at the Marine Biological Laboratory (MBL) for their support; the National Cancer Institute electron microscopy core; the NHLBI light microscopy, bioinformatics, DNA sequencing and genomics, flow cytometry and animal facility core facilities.

## Materials and Methods

### Cell and Organoid Culture

1205Lu cells (a kind gift from Dr. Glenn Merlino, NCI) were cultured in Tu 2% media (80% MCDB153, 20% L-15) supplemented with 2% fetal bovine serum (FBS; Atlanta Biologicals S11150) 2.5 ng/ml insulin (Sigma), and 1.68mm CaCl_2_. Immortalized human melanocytes (Dr. Glenn Merlino, NCI) were cultured in dermal cell basal media (ATCC, PCS-200-030), supplemented with melanocyte growth kit (ATCC, PCS-200-041). MCF10A (ATCC, CRL-10317), PC-3 (ATCC, CRL-1435), HEK293FT (ATCC, CRL-3216) and T47D (NIH, NCI-60) cells were all cultured in DMEM/F12 + GlutaMax (Gibco) with 10% fetal bovine serum (Atlanta Biologicals), and 1% penicillin and streptomycin (ThermoFisher, 10378016). RWPE-1 cells were cultured in keratinocyte serum-free media supplemented with 25 mg of bovine pituitary extract and 2.5 µg epidermal growth factor (ThermoFisher, 17005042). All cells were maintained at 37°C at 5% CO_2_ for <15 passages. Transient expression of cDNAs or siRNAs was performed using the Amaxa nucleofector kit R (Lonza) and Amaxa Nucleofector II (Lonza) programs X-001 (1205Lu), U-024 (melanocytes), or the Amaxa nuclefector kit V (Lonza) and Amaxa Nucleofector II (Lonza) programs T-020 (MCF10A), T-013(PC-3, RWPE-1), Q-001 (Hek293FT), X-005 (T47D) and 1.5 µg of DNA for 2×10^6^ cells.

To generate melanoma organoids, 2000 cells were seeded in 200 µl of media in low adherence U-bottom 96 well dishes (Nunclon Sphera, Thermo Fisher) and allowed to proliferate for 7 days. To embed organoids in ECM, 3 thin layers of polymerized collagen were used to control Z height and placement of organoids in a cell culture dish. Briefly, 35mm glass bottom dishes (FluoroDish, WPI) were plasma cleaned then coated with a thin layer of 2.5 mg/ml Type 1 collagen (Corning) which was allowed to polymerize at 37°C for 1 hour. Following polymerization, organoids were gently aspirated from 96 well dishes using a pipette and individually placed in the center of the 35mm dish and encased with an additional layer of 2.5 mg/ml of collagen that was allowed to polymerize at 37°C for 1 hour. Following the second layer, a final layer of collagen was applied to ensure full coverage of the organoids and attachment of the collagen to the dish sidewalls. It was treated similarly to the other layers. Upon final polymerization, 1 ml of culture media was applied to each dish and organoids were maintained for 1 week to allow for proliferation and cell invasion into the ECM.

### CDNA expression vectors and lentiviral expression

The following cDNAs were used for DNA transfections or subcloning: mCherry NLS, mCherry-N1, and mEmerald-H2B were kind gifts of the late Mike Davidson (Florida State University, Tallahassee, FL); GFP-cGas (a kind gift from Dr. Hawa Racine Thiam, Stanford) ;GFP-Lap2b (Addgene 62044); EGFP-LBR (Addgene 128150); pTrip-CMV-GFP-Flag-cGas (Addgene 86675); pLBS.CAG-NLS-mScarlet (Addgene 129336). To generate mCherry-LBR, LBR cDNA was PCR amplified to generate NheI and EcoRI restriction sites and an 18 amino acid linker using primers (Fw-CGT TCC GCT AGC GCC ACC ATG CCA AGT AGG AAA TTT GCC GAT GGT GAA GTG GTA AGA GGT C) (Rev-CCA GAC GAA TTC CGC TGT AGA TGT ATG GAA ATA TAC GGT AGG GCA CAC GCT GAC AGT AC). To generate mCherry-LBR-ΔTudor+RS, the reverse primer above was reused and the following forward primers was used (CGT TCC GCT AGC GCC ACC ATG GGT CGA CCA CCT AAA AGT GCC CGC CGA TCT. The obtained inserts and mCherry-N1 were digested with restriction enzymes (NheI and EcoRI, New England Biolabs) then ligated with T4 DNA ligase (ThermoFisher, EL001) and treated with calf intestinal alkaline phosphatase (CIAP, ThermoFisher, 18009019) and gel purified (QIAquick Gel Extraction Kit; Qiagen) and transformed into DH5α competent cells (ThermoFisher, 18258012). To generate mCherry-LBR-R583Q and m-Cherry-LBR-Y468T, site directed mutagenesis was used (QuikChange II XL Site Directed Mutagenesis Kit, Agilent, 200521) as directed. The primers used to introduce the appropriate mutations into mCherry-LBR-N-18 were (5’-ctcgtcacgagcttcttggtggacaagcaacat-3’), (5’-atgttgcttgtccaccaagaagctcgtgacgag-3’) for R583Q and (5’cttggaagctgtaataaagggaacccacaccaagt-3’), (5’-acttggtgtgggttccctttattacagcttccaag-3’) for Y468T. All colonies were then miniprepped (Qiagen) and confirmed by sequencing, then large scale purified (Qiagen) for experimental use. For KD of candidate NE proteins for screening for their effects on nuclear fragility in confinement, ON-TARGETplus SMARTpool siRNAs (Dharmacon) were used for human LBR (J-021505-05), emerin (J-011025-06) Nup93(J-020767-09), Lap2β (J-027195-09), CHMP2A (L-020247-01), laminB2 (L-005290), laminA (L-004978), and non-targeting pool (D-001810-10). For LBR rescue experiments, a custom 5’UTR duplex was utilized (5’GUAAAGGAGUGCUGUCUUAUU 3’) (Dharmacon). For all siRNA experiments, cells were transiently transfected with 1 µM of siRNA for 48 hours prior to experimental use.

Generation of a custom LBR shRNA lentivirus plasmid was done using the above 5’UTR duplex RNA sequence to design a shRNA ligated into the pLKO-Tet-On lentivirus AgeI-EcoRI sites for expression under the control of the H1/TO promoter (*56*). As a control, inducible NT expressing lentivirus was used (Dharmacon). 1205Lu cells stably expressing LBR shRNA (Dharmacon) and NT shRNA (Dharmacon) were generated using the second generation lentiviral packaging plasmid psPax2 (Addgene 12260) and pMD2.G (Addgene 12259). Lentiviral particles were generated by first transfecting 1 million HEK293FT with Lipofectamine 2000 (Thermo Fisher) according to manufacturer’s protocols using 2.5 µg plasmid DNA, 1 µg pMD2.G, and 2.5 µg psPax2 and collecting the virus containing supernatant at 48 hours. Supernatant was then filtered through a 0.45 µm filter (Sigma), then immediately placed on target 1205Lu cells that had been seeded 24 hours before transduction at a 1:1 ratio of viral supernatant to 2% Tu media and allowed to incubate for 48 hours, upon which cells were passaged into media containing a final concentration of 2.0 µg/ml puromycin for selection. Knockdown was induced in cell culture using 100 nM doxycycline (Sigma) for 48 hours prior to experiments.

### Generation of stable cell lines

1205Lu cells stably expressing pTrip-CMV-GFP-Flag-cGas (Addgene 86675) and pLBS.CAG-NLS-mScarlet (Addgene 129336) were generated using the second generation lentiviral packaging plasmid psPax2 (Addgene 12260) and pMD2.G (Addgene 12259) as described above. Following 2.0 µg/ml puromycin selection, cells were then expanded for 2 weeks before being sorted by flow cytometry for GFP and mScarlet positive cells.

### *Ex vivo* tumor models

Tumors were generated by intradermal injection of 5000 human metastatic melanoma cells (1205Lu) in anaesthetized Crl:NU(NCr)-Foxn1nu (athymic nude mice; Charles River, Rockville MD). Cells stably expressing GFP-cGas and mScarlet NLS were injected in a 1:1 mixture of Matrigel and minimal essential medium (Gibco) and imaged once tumors were between 4-5 mm in size. For tumor imaging, animals were euthanized and dorsal skin was removed with at least 10 mm of margin around the tumor. The superficial fat layer was removed and tumors were treated with Hoechst 33342 (1:1000) for 10 min at RT, then whole-mounted tumors *in situ* were imaged in dermal cell basal media supplemented with melanocyte growth kit. Animal protocols and care were performed as directed by the National Heart Lung and Blood Institute and approved by the Institutional Animal Care and Use Committees guidelines.

### Drug Treatments

The following pharmacological inhibitors were used:

Simvastatin: (5µM, 8 µM, 20 µM, dissolved in DMSO; Sigma).

### RNA-Sequencing

RNA from 1205Lu cells and immortalized human melanocytes were isolated and purified using RNeasy Micro Kit (Qiagen) according to manufacturer’s instruction and concentration was quantified using a NanoDrop Spectrophotometer (ThermoScientific). Sequencing libraries were constructed from 100– 500 ng of total RNA using Illumina’s TruSeq Stranded Total RNA Kit with Ribo-Zero according to manufacturer’s instructions. The fragment size of RNAseq libraries was verified using a 2100 Bioanalyzer (Agilent) and the concentrations were determined using a Qubit instrument (LifeTech). The libraries were first run on an Illumina Miseqkit for QC purposes. After QC analysis, the libraries were loaded onto an Illumina HiSeq 3000 for 2 x 75-bp paired end read sequencing. The fastq files were generated using the bcl2fastq software for further analysis.

### Bioinformatics

#### The Cancer Genome Atlas (TCGA) Data Acquisition

Unbiased identification of potential nuclear membrane genes of interest was obtained using the uniform manifold approximation and projection (*57*) (UMAP) subcellular distribution analysis from The Human Protein Atlas. Querying the ‘Nuclear Membrane’ category resulted in the identification of 249 nuclear membrane associated genes. Next, The Cancer Genome Atlas (TCGA) Pan-Cancer dataset was accessed through the University of California, Santa Cruz (UCSC) Xena Browser (http://xena.ucsc.edu/), and was queried for 10 cancer subtypes; Bladder Urothelial Carcinoma (BLCA), Breast invasive carcinoma (BRCA), Colon adenocarcinoma (COAD), Head and Neck squamous cell carcinoma (HNSC), Kidney renal papillary cell carcinoma (KIRP), Liver hepatocellular carcinoma (LIHC), Lung adenocarcinoma (LUAD), Lung squamous cell carcinoma (LUSC), Prostate adenocarcinoma (PRAD) and Stomach adenocarcinoma (STAD). The subset of 249 NE genes identified from the Human Protein Atlas was surveyed in the Xena Browser then exported for subsequent data visualization with Prism.

#### Identification of genes of interest from patient melanoma samples

Patient bulk melanoma tumor RNA-seq data was downloaded from GEO (GSE98394) and categorized based on provided clinical staging information into benign nevi (27 samples), stage 1 and 2 (35 samples), and stage 3 and 4 (32 samples)(*28*).Clinical staging information was based on both the Tumor, Nodes, Metastasis classification of malignant tumors (TNM) and American Joint Committee on Cancer (AJCC). Briefly, benign nevi and stage 1 tumors were <1 mm in thickness, and presented with no signs of cancer in the lymph nodes, and were termed “non-invasive”. While stage 4 represented patients with tumors >4 mm in thickness and lymph nodes positive for cancer were termed “invasive”. Melanoma single cell RNA-seq (scRNA-seq) data was downloaded from GEO (GSE72056) and malignant samples were isolated (*31*). Melanoma cell line RNA-seq was downloaded from the Cancer Cell Line Encyclopedia (CCLE) and malignant melanoma samples were isolated. All RNA-seq data was subjected to quality control analysis using FastQC (v0.11.8)(*58*). This was followed by alignment to the human genome (GRCh38/hg38) using spliced transcripts aligned to a reference (STAR)(*59*). Gene expression was quantified using featureCounts (v2.0.0)(*60*). RNAseq sample quality was determined using ESTIMATE (v 1.0.13) R package. Tumor purity estimates and sex were used as the covariates for differential expression analysis, performed using limma-voom (*61*). Genes with significantly differential expression with p value < 0.05 were subjected to pathway enrichment analysis using clusterProfiler (v3.14.3)(*62*). All RNA-seq data was filtered to remove genes with an expression count < 5 in 90% of the samples, and FDR< 0.05, to remove low-expressing and non-significant genes. Integration of data sets was performed in the following order: bulk tumor (GEO: GSE98394), single cell melanoma samples (GEO: GSE72056), and Broad CCLE melanoma cell lines. Genes were retained if they were present in all datasets, and consistently up- or down-regulated between all samples, forming “Group 1”. Following this, differential expression analysis (DEA) was performed as previously described between 1205Lu and immortalized human melanocyte (IM) cells, to identify transcriptomic changes occurring in 1205Lu as compared to IM. The DEA was then integrated with the previously described Group 1 using the same criteria as listed above. The final gene set was then named “Group 2” and used for analysis. To determine if any gene ontology (GO) terms were selectively enriched, the gene set from Group 2 was input into the GO cellular components (P < 0.05 was considered significant), resulting in the identification of the ‘nucleus’ GO term (GO:0005634) as the second most enriched category. Additional ranking of genes by q-value and fold change, and targeted segregating of nuclear genes that have a known role in nuclear envelope mechanics resulted in our final list of gene candidates for further analysis.

### Confinement assays

To reproducibly confine cells during high-resolution live-cell imaging, confinement was performed using the 1-well Dynamic Cell Confiner System (4Dcell) per manufacturer’s directions. Briefly, the system consists of a cobalt autonomous vacuum pump attached to a manufactured PDMS suction cup fitted with a glass coverslip containing PDMS micropillars of 2.5, 3, or 5 µm height which determine the spacing between the top of the coverslip and the cell culture dish. To perform confinement experiments, cells were transiently transfected and plated for 24-48 hours (depending on experimental conditions) on 35mm glass bottom dishes (FluoroDish, WPI) coated with 10 µg/ml fibronectin. Prior to imaging, the PDMS suction cup and coverslip were briefly sonicated in 70% ethanol, followed by a 5-minute sonication in PBS (Invitrogen), then placed in cell culture media for 1 hour at room temp to equilibrate the PDMS. To confine the cells, the 35mm glass bottom dish was washed briefly with pre-warmed PBS, then 1 ml of fresh warmed cell culture media was added. The sample was then placed on the microscope and the PDMS suction cup was attached by a low pressure vacuum seal that was sufficient for attachment of the PDMS perimeter base to the glass dish, but insufficient for cellular confinement. Confinement was initiated over 1 minute using a ramp of 10 mbar per second utilizing the custom software (4Dcell) to control the vacuum pump. After cells were confined, images were acquired from multiple (10–40) XY positions to ensure full unbiased coverage of the confined area, images were acquired every 1-2 minutes over 1 hour for analysis.

### Immunofluorescence

#### Immunostaining of adherent cells

Cellular immunofluorescence for nuclear envelope proteins was performed using 1205Lu cells plated on cleaned coverslips (#1.5, 22 mm x 22 mm, Corning) for 24-72 hours depending on experimental conditions. Samples were then fixed for 20 min at 37°C with 4% paraformaldehyde (PFA; Electron Microscopy Science,15710) in cytoskeleton buffer (CB; 10 mM MES, 3 mM MgCl2, 138 mM KCl, 2 mM EGTA), permeabilized with 0.5% Triton X-100 in CB at RT for 5 mins, and quenched with 10 mM glycine in CB at RT. Cells were washed 2 x 5 mins then 2 x 10 mins with Tris Buffered Saline (TBS; 20 mM Tris, pH 7.6, 137 mM NaCl2) before blocking for 1 hour at RT with blocking solution (2% BSA IgG free and protease free (Sigma-Aldrich, A3059); 0.1 % Tween 20 (Sigma-Aldrich, P1379) in TBS). Cells were incubated for 1.5 hours at RT with the following primary antibodies: LBR (1:200, Atlas Antibodies HPA062236), LaminB1 (1:500, Abcam ab16048), LaminB2 (1:500, Abcam ab8983), Sun2 (1:500, Abcam ab87036), H4K20me2 (1:500, Abcam ab9052), H3K9me2 (1:500, Active Motif AB 2793199), Lap2 (1:1000, BD Transduction Laboratories 611000), Mab414 (1:1000, Abcam ab24609), Lamin A/C (1:500, Abcam ab108595) or HP1 (1:500, Abcam ab109028), diluted in blocking solution then washed in TBS 3 x10 mins before being incubated with fluorophore-conjugated secondary antibodies (1:500), Alexa Fluor 488 Donkey anti-rabbit (711-545-5152), Alexa Fluor 594 Donkey anti-rabbit (711-585-152), Alexa Fluor 488 Donkey anti-mouse (715-545-150) and Alexa Fluor 594 Donkey anti-mouse (715-585-150), Alexa-Fluor-647 phalloidin (1:200; ThermoFisher, A22287) and Dapi (1µg/mL; Sigma, 268298) diluted in blocking solution, for 1 hour at RT. Cells were washed with blocking solution with TBS (2 x 10 mins, each). Coverslips were mounted on glass slides in mounting media (Dako; Pathology Products, S3023).

#### Immunostaining of confined cells

Immunostaining for laminA in confined 1205Lu cells was performed in cells plated on 35mm glass bottom dishes (FluoroDish, WPI) and confined to 3µm. Samples were then fixed during confinement for 60 min at 37°C with 4% paraformaldehyde (PFA; Electron Microscopy Science,15710) in CB to allow fixative to penetrate the PDMS cell confiner. Following fixation, the cell confiner was removed and dishes were washed 2x 5 min with TBS to remove cell debris then treated as described above.

#### Immunostaining of organoids

Tumor organoids embedded in collagen in 35mm glass-bottom dishes were fixed for 60 min at 37°C with 4% paraformaldehyde in CB then permeabilized with 0.5% of Triton X-100 in CB at 4°C overnight. Organoids were then washed 4 x 15 mins with TBS before blocking at 4°C overnight. Organoids were then incubated at 4°C overnight with Dapi (1µg/mL; Sigma, 268298) and Alexa-Fluor-647 phalloidin (1:50; ThermoFisher, A22287) diluted in blocking solution. Organoids were washed with blocking solution then with TBS (2 x 10 mins, each).

#### Immunostaining of tissue microarrays

Tissue microarrays of melanoma progression (ME1004f, US Biomax) were used for visualization and quantification of nuclear morphology alterations during cancer progression. Paraffin was removed through a series of xylene and ethanol dilution rinses: 100% xylene, 3 x 5 min, 100% ethanol, 2 x 5 min, 95% ethanol, 1 x 5 min, 70% ethanol, 1 x 5 min, PBS 2 x 5 min, 1 x 10 min PBS + 0.05% Tween 20. Antigen retrieval was performed in sodium citrate buffer (10mM sodium citrate in PBS/0.05% Tween 20) using a microwave to maintain 95°C for 25 min, then TMAs were blocked with 2% BSA at 4°C overnight. Following blocking, TMAs were transferred to humidity chambers and incubated for 2 hrs at RT with Alexa-Fluor-488 Wheat Germ Agglutinin (1:100; ThermoFisher, W11261) and Dapi (1µg/mL; Sigma, 268298) diluted in blocking solution, for 1 hour at RT. Cells were then washed in PBS 5 x 5 mins, then mounted with a cleaned glass coverslip (#1.5, 24 mm x 60 mm) in mounting media (Dako; Pathology Products, S3023).

### Microscopy

#### Spinning Disk Confocal Imaging

Imaging was performed on a Nikon Eclipse Ti2 microscope equipped with a Yokogawa CSU-W1 spinning disk scanhead, a Nikon motorized stage with a Nano-Z100 piezo insert (Mad City, Madison, WI), and either a Plan Apo 60x oil 1.49 NA DIC, SR Plan Apo 60x 1.2 NA water objective or an Apo TIRF 100x oil 1.49 NA DIC Nikon objective lens. Illumination was provided by a Nikon LUNV 6-line laser unit, and images were captured with a Hamamatsu Orca-Flash 4.0 v3 camera. The system was controlled by NIS-Elements software (Nikon). Cells were imaged either on coverslips (#1.5, 22 mm x 22 mm) or in 35 mm glass bottom dishes (FluoroDish) depending on experimental conditions. For the siRNA-based confinement screen, imaging was also performed on a Nikon Eclipse Ti microscope equipped with a Yokogawa CSU-X1 spinning disc scanhead, a Hamamatsu Orca-flash 4.0 v2 camera and a Plan Apo 60x oil 1.4 NA objective. Illumination was provided by a 4-line Agilent MLC400B Monolithic Laser Combiner. The microscope was similarly equipped with a Nano-Z100 piezo insert (Mad City) and controlled by NIS-Elements software (Nikon). For live cell imaging, cells were incubated with Hoechst 33342 (1:1000; ThermoFisher), ER-Tracker blue-white (30 min at 1 µM; ThermoFisher), or SirDNA (1 µM; Cytoskeleton) at 37°C at 5% CO_2_, then washed and resuspended in 1 ml of cell culture media. Maintenance of sample temperature and humidity was performed by a Tokai Hit stage-top incubator (Tokai Hit). To image tumor organoids, cell culture dishes were imaged by spinning disc confocal microscopy with a SR Plan Apo 60x 1.2 NA water objective (Nikon). For each organoid, an 8×8 tile was used to capture the full organoid and area of cell invasion and a z-stack was taken at 0.3 µm intervals for each example for later image analysis.

#### Super-resolution Confocal Imaging

Imaging of confined live cells for superresolution was performed using a LSM 880 Zeiss confocal microscope equipped with an Airyscan using a Plan-Apo 63x 1.4 NA oil objective. Airyscan image reconstructions were processed in auto strength mode using ZenBlack software (Version 2.3). Additional analysis was performed in ImageJ (NIH).

#### Two-Photon imaging

Two-photon imaging of whole-mounted 1205Lu melanoma tumors that had been resected from mice was performed using an upright Leica SP8 DIVE (Deep In Vivo Explorer) 2-photon system using a Leica HC Fluotar L VISIR 25x/0.95 Water immersion objective lens (working distance 2.4 mm) and equipped with a FALCON FLIM module (Leica Microsystems, Mannheim, Germany). For 2-photon excitation, an infra-red laser, InSight X3 (Spectra-Physics), dual-beam with one fixed 1045-nm line and one tunable line from 680 to 1300 nm was used. Sequential excitation at 1045 nm (InSight laser) was used for Hoechst, second harmonic generated intrinsic signal (SHG) from fibrillar collagen, and mScarlet, while a second imaging sequence with an excitation of 950 nm was used for EGFP fluorescence. Acquisition detection was on 4 spectrally tunable HyD detectors (NDDs) and their range was set as follows HyD-NDD1 (420–465 nm) for Hoechst, HyD-NDD2 (517–528 nm) for SHG, HyD-NDD3 (530–560 nm) for EGFP, and HyD-NDD4 (666–685 nm) for mScarlet, respectively.

#### Transmission Electron Microscopy (TEM)

*In situ* fixation of cell monolayers on a glass coverslip for TEM was performed using an EM fixative solution (2% glutaraldehyde in 0.1M cacodylate buffer) for 1hr at room temperature, then stored at 4^0^C. Sample processing was performed in Cacodylate buffer (0.1M, pH7.4), and fixed in osmium (1% in 0.1m Cacodylate buffer) for 1hr at room temperature, then dehydrated in a series of alcohol dilutions, infiltrated with epoxy resin (Polyscience) and embedded. Resin blocks were thin sectioned (70 nm) and mounted on 200 mesh copper grids and counterstained with aqueous uranyl acetate (0.5%) and Reynolds lead citrate. Images were acquired with a Hitachi H7600 TEM.

### Image analysis

#### Quantification of NE permeability and integrity during confinement

For all PDMS confinement assays, time-lapse spinning disk confocal movies of cells transiently transfected with relevant plasmids were used to quantify the percentage of events in the following conditions: Nuclear envelope rupture, defined as the appearance of a perinuclear GFP-cGas foci post confinement. Nuclear envelope permeability, defined as the leakage of the mCherry-NLS signal from the nucleus into the cytosol of the cell post confinement; Measurement of INM and ONM rupture as defined by formation of a discontinuity of the INM marker Lap2β and the ONM marker ER-tracker, respectively. For all conditions, only adherent cells labeled with all relevant fluorescent markers and which remained in the view field both pre- and post-confinement were included in the analysis. Additionally, any cells with a ruptured or permeable NE prior to confinement, or undergoing division, were not included in the quantification. For analysis of NE permeability (mCherry-NLS leakage) and NE rupture (GFP-cGas foci), 10-40 imaging fields were randomly chosen per dish to ensure unbiased coverage of the confined area. To determine a baseline NE integrity percentage, the first 2 time points prior to initiation of cellular confinement, as described above, were analyzed and quantified for total intact nuclei. Following initiation of confinement, images were acquired every 1-2 min for a standard duration of 60 min and quantified again for total intact or ruptured nuclei using the last 2 time points. All movies were analyzed individually and scored based on the total percentage of mCherry-NLS and GFP-cGas positive cells from the sum of all fields of view in each cell culture dish. Any dish containing less than 200 cells was excluded to minimize skewing of data due to low cell density.

For analysis of nuclear integrity in 3D tumor organoids, organoids were generated, imaged and fixed and stained as described above and analyzed using Imaris v9 (Bitplane, Switzerland). First, Imaris surface rendering utilizing the phalloidin channel was used to segment the whole tumor organoid and to hand segment the core of the tumor organoid, defined as the center of the organoid approximately 200-250 µm from tumor edge. The centroid of this area was used to quantify cell migration distance from the tumor core. Next, using Imaris spot detection and the DAPI channel, individual nuclei were counted in the tumor spheroid edge (TE) and invasion zone (IZ) and classified as either positive or negative for NE integrity based on the presence or absence of a perinuclear GFP-cGas signal that colocalized with the DAPI channel. Each organoid was given a percentage score of GFP-cGas foci (cGas positive nuclei/ total nuclei) in each organoid region.

For *ex vivo* tumor analysis, we collected a series of x-y-z images (typically 1 × 1 × 3 μm^3^ voxel size) along the *z*-axis at 3 μm intervals over a 5×4 mm region with a range of 100–220 μm z depth throughout the whole-mount tumors, over using the tile function (Navigator) of the Leica LAS-X software to automatically generate stitched volumes. For 3D renderings, segmentations and quantitative image analyses, we used Imaris v 9 or 10.0.1 software.

#### Nuclear Height

Quantification of nuclear height pre- and post-confinement was performed using a spinning disk confocal and a SR Plan Apo 60x 1.2 NA water objective (Nikon). To determine nuclear height, Z-stacks were taken every 0.3 µm pre and post cellular confinement for 3D image reconstruction. Hand-drawn line ROIs were generated using ImageJ (NIH) to measure the approximate Z height of the nucleus.

#### Nuclear Envelope Width

Analysis of NE spacing was performed using TEM images of immortalized melanocytes (14 cells) and 1205Lu cells (11 cells) using hand drawn ROI’s measuring the distance from the INM and ONM over a distance of 200nm at the nuclear periphery. Briefly, using ImageJ (NIH) a 1 pixel line was drawn from the membrane boundary of the INM to the membrane boundary of the ONM when both membranes could be clearly resolved in the image, over a distance of 200nm, resulting in 50-100 measurements per cell.

#### Nuclear Morphology from Tumor Microarrays and in 3D

Analysis of nuclear morphology of cells in tumor microarrays of melanoma cancer progression (ME1004F, Biomax) was performed using 5 fields of view taken from each patient sample representing benign nevi (13 samples), stage 1 and 2 (23 samples), stage 3 (4 samples) and lymph node metastasis (11 samples). After immunofluorescence was performed, a custom made ImageJ macro for automatic segmentation of the nucleus based on the DAPI channel using Otsu segmentation, was applied to the images to quantify NE solidity (Area/Convex Area).

#### Bleb perimeter and area

Quantification of NE bleb morphology was performed from time-lapse movies of cells transiently transfected with GFP-cGas and mCherry NLS. To determine maximum bleb perimeter and area, cells exhibiting NE blebs representing 1205Lu WT (116 cells), and LBR KD (139) were isolated to generate individual fields of view. Using the mCherry-NLS channel for segmentation, ROI’s were hand drawn around the frame representing the maximum bleb perimeter and quantified for shape descriptors.

#### Analysis of organoid and tumor volume

Analysis of 3D organoid and tumors was performed using samples generated and imaged as described above. For organoids, segmentation was performed utilizing the phalloidin channel, for *ex vivo* tumors the Hoechst channel. In both samples only cells in tumor bulk not invading into the ECM were used to calculate 3D volume. Data was then exported to excel and transferred to Prism for statistical analysis.

### Atomic force microscopy (AFM)

Immortalized human melanocytes (IM) or melanoma line 1205Lu cells expressing either GFP-H2B or GFP-Lap2β (to visualize the nucleus) were plated on glass-bottom dishes (Willco Wells) coated with 10µg/mL of fibronectin. AFM force spectroscopy experiments were performed using a Bruker BioScope Catalyst AFM system mounted on an inverted Axiovert 200M microscope (Zeiss) equipped with a confocal laser scanning microscope 510 Meta (Zeiss) and a 40x objective lens (0.95 NA, Plan-Apochromat, Zeiss). The microscope system was placed on an acoustic isolation table (Kinetic Systems). During AFM experiments cells were maintained at 37°C using a heated stage (Bruker). A modified AFM microcantilever with an attached 25 µm polystyrene bead (Novascan) was used for all AFM measurements. All AFM microcantilevers were pre-calibrated using the standard thermal noise fluctuations calibration method. The calibrated spring constants were 0.6N/m-1.2N/m. For nuclear measurements, five force curves were performed in succession with a 30 sec delay between each measurement. The applied force was set to be between 20nN – 40nN yielding indentations between 1.5µm-3µm, allowing us to press against and deform the cell’s nuclei, confirmed by confocal microscopy. The force curves ramp rate was set to 0.5Hz yielding AFM probe approach/compressive speeds between 3µm/s-5µm/s. The nucleus effective stiffness (N/m) was analyzed and determined using the Bruker NanoScope Analysis software. In brief, force curves were corrected for the non-contact region slope (typically arising from the hydrodynamic drag and AFM probe-sample orientation) using a baseline function. To separate the nuclear mechanical component from the cortical component, we first utilized sharp AFM probes (PFQNM-LC, Bruker) to determine that the nuclear depth in melanocytes and 1205 Lu cells was approximately 1.5µm in depth after penetrating the plasma membrane as indicated by the change in stiffness on a force curve. Due to this, we selected a region in the force curve ∼1.5µm beyond the initial indentation where we then fit a linear regression to extract the nuclear effective stiffness for all experimental conditions.

### Measurement of cellular cholesterol

Measurement of bulk cellular cholesterol was performed using the Amplex Red Cholesterol Assay (Molecular Probes, Thermo Fisher Scientific) per manufactures instructions. Briefly, cells were cultured for 7 days in Tu 2% media (1205Lu) supplemented with lipid depleted FBS (Omega Scientific, Fisher Scientific). Dox inducible LBR KD was initiated 3 days prior to the experiment, followed by expression of LBR constructs 24 hours prior to analysis. Directly before the assay, 2000 cells were seeded into each well of a black walled 96 well imaging dish (Thermo Fisher) and incubated for 30 min at 37°C in the presence of 2U/mL HRP, 2 U/mL cholesterol oxidase, 0.2 U/ml cholesterol esterase and 300µM Amplex red reagent. Fluorescence was measured using the iBright 1500 (Thermo Fisher) and quantified using ImageJ. Intensity measurements were determined by first subtracting background fluorescence from a no-cholesterol control well and normalizing each sample to the 1205Lu WT condition. To measure local variation in relative cholesterol concentration within individual cells, cells were stained with Bodipy cholesterol (TopFluor-Cholesterol, Avanti Polar Lipids) and analyzed by superresolution confocal microscopy. To generate labeling solution, Bodipy cholesterol was complexed with methyl-β-cyclodextrin (MβCD, Sigma). First Bodipy-Cholesterol was dissolved in a 1:1 (v:v) solution of chloroform and methanol to a final concentration of 1 mg/ml in a glass tube, then dried with a stream of nitrogen gas, followed by the addition of 370 mM MβCD in PBS at a 100:1 ratio. Solution was then votexed, sonicated in a water bath for 3 minutes, and incubated on a rocker overnight at 37°C. Undissolved dye was then removed by centrifugation at 16,000 x g for 15 min. 1205 Lu cells with or without molecular perturbations of LBR were incubated with the Bodipy-cholesterol/MβCD solution at 1:2000 dilution for 10 minutes. Cells were then incubated in fresh media containing Blue-White DPX ER tracker (to label the endoplasmic reticulum (RE), 1 µM; Thermo) and SiR-DNA (to label the nucleus, 1 µM; Cytoskeleton) for 1 hour to allow for cholesterol internalization to the cell membranes and uptake of vital dyes. Cells were then subjected to times lapse superresolution microscopy to visualize labeled cholesterol, the ER, and nucleus. Analysis of local and total cholesterol levels was performed in ImageJ. Briefly, supreresolution Z-stack images representing 1205Lu WT (18 cells) and LBR KD (18 cells) were selected and the middle z-plane was isolated for analysis. Using ImageJ the entire cell was thresholded by cholesterol intensity, background subtracted, and quantified for total cholesterol intensity in a single z slice. To assess local enrichment of cholesterol, images containing nuclear blebs in 1205Lu WT (10 cells) and LBR KD (9 cells) were selected and two measurements were taken using a 2µm long line scan 5 pixels in width at the bleb neck, nuclear envelope, and the ER. All intensities were background subtracted, and normalized to the total cholesterol intensity in that z plane for each cell. Paired points representing each area were then quantified for statistical significance. To inhibit production of endogenous cholesterol, cells were treated with either 5µM or 20µM simvastatin dissolved in DMSO for 48 hours prior to any experimental analysis.

### Western blot analysis

Cells were lysed in Laemmli sample buffer, separated by SDS-PAGE and electro-transferred at room temp (RT) for 1 hour to Immobilon-P PVDF membrane. Membranes were blocked for 1 hour at RT with 5% nonfat dry milk (wt/vol) in TBS-T buffer (TBS + 0.1% Tween-20 [vol/vol]), incubated for 2 h at RT with indicated primary antibodies, washed 3 x 5 min in TBS-T, incubated with appropriate HRP-conjugated secondary antibodies (1:10,000) for 1h at RT, then washed 3 x 5 min in TBS-T. An ECL detection system (Millipore) was used to visualize protein bands. The following antibodies were used: anti-LBR (1:500; Abcam ab32535), anti-lamin A/C (1:1000; Abcam, ab108595), rabbit anti-lamin B1 (1:1000; Abcam, ab16048), mouse anti-lamin B2 (1:1000; Abcam, ab8983), rabbit anti-GAPDH (1:2000; Clone 14C10, Cell Signaling, 2118S). The secondary antibodies (HRP-conjugated goat anti-mouse (115-035-003) or goat anti-rabbit (111-035-003)) were from Jackson ImmunoResearch Laboratories. Quantitative analysis was achieved by first performing local background subtraction around bands of interest, then calculating the ratio between the band of interest and the loading control (GAPDH). For analysis of protein levels relative to WT conditions, intensities were further normalized to WT levels to obtain percent relative to WT using ImageJ (NIH).

### Statistical analysis

Statistical analysis was performed in Prism (GraphPad Software version 9). Graphs represent mean values± SEM or ± SD as indicated in the figure legends of at least three independent experiments. Statistical significance was obtained using ANOVA or student t-test with Welch’s, Kolmogorov-Smirnov or Mann-Whitney correction as indicated in the figure legends. Significance is reported as *p ≤ 0.05, ** p ≤ 0.01, ***p ≤ 0.001, and ****p ≤ 0.0001.

---

**Fig. S1.**
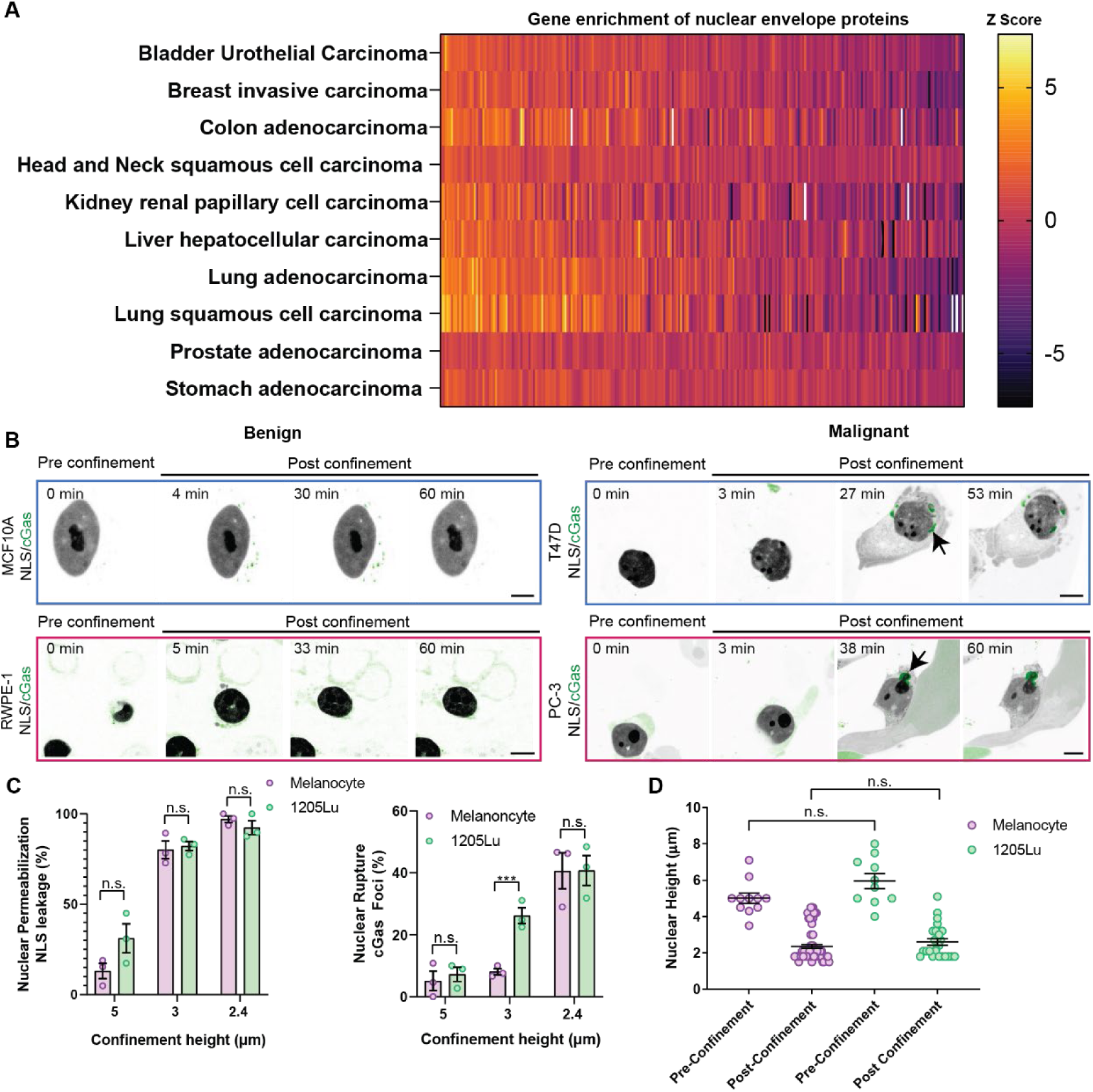
**(A)** Heat map of transcript abundance (normalized relative to a baseline human genome) of 249 genes encoding nuclear envelope proteins (curated from the Human Protein Atlas) from RNA-seq data from 6860 clinical samples representing 10 cancer subtypes from The Cancer Genome Atlas (TCGA) PanCancer repository. **(B)** Confocal image series of living MCF10A (upper left) or T47D (upper right) breast cells (blue outline) or RWPE-1 (lower left) or PC-3 (lower right) prostate cells (pink outline) transfected with mCherry-NLS (inverted grayscale) and GFP-cGas (green) before and after confinement to 3µm. Arrows highlight cGas foci. Bar = 10µm. **(C)** Quantification of NE permeabilization (left) and NE rupture (right) in human melanocytes (purple) and 1205Lu cells (green) at different confinement heights (Melanocytes n=3 experiments for all heights, n=404 cells (2.4µm), n=196 cells (3µm), n=184 cells (5µm); 1205Lu n=3 experiments for all heights, n=301 cells (2.4µm) n=604 cells (3µm), n=1375 cells (5µm). **(D)** Nuclear height of fixed 1205Lu cells (green) and human melanocytes (purple) expressing GFP-NLS measured from confocal image series before and after confinement to 3µm. 1205Lu: before confinement n=3 experiments, 189 cells; after confinement n=3 experiments, 159 cells. Human melanocytes: before confinement n=3 experiments, 107 cells; after confinement n=3 experiments, 71 cells. Data points represent independent experiments each consisting of pooled cell data from a single imaging dish (C) or individual cells (D). Significance was tested with a Student’s T-test Mann-Whitney, bar= mean, error bars= SEM.

**Fig. S2.**
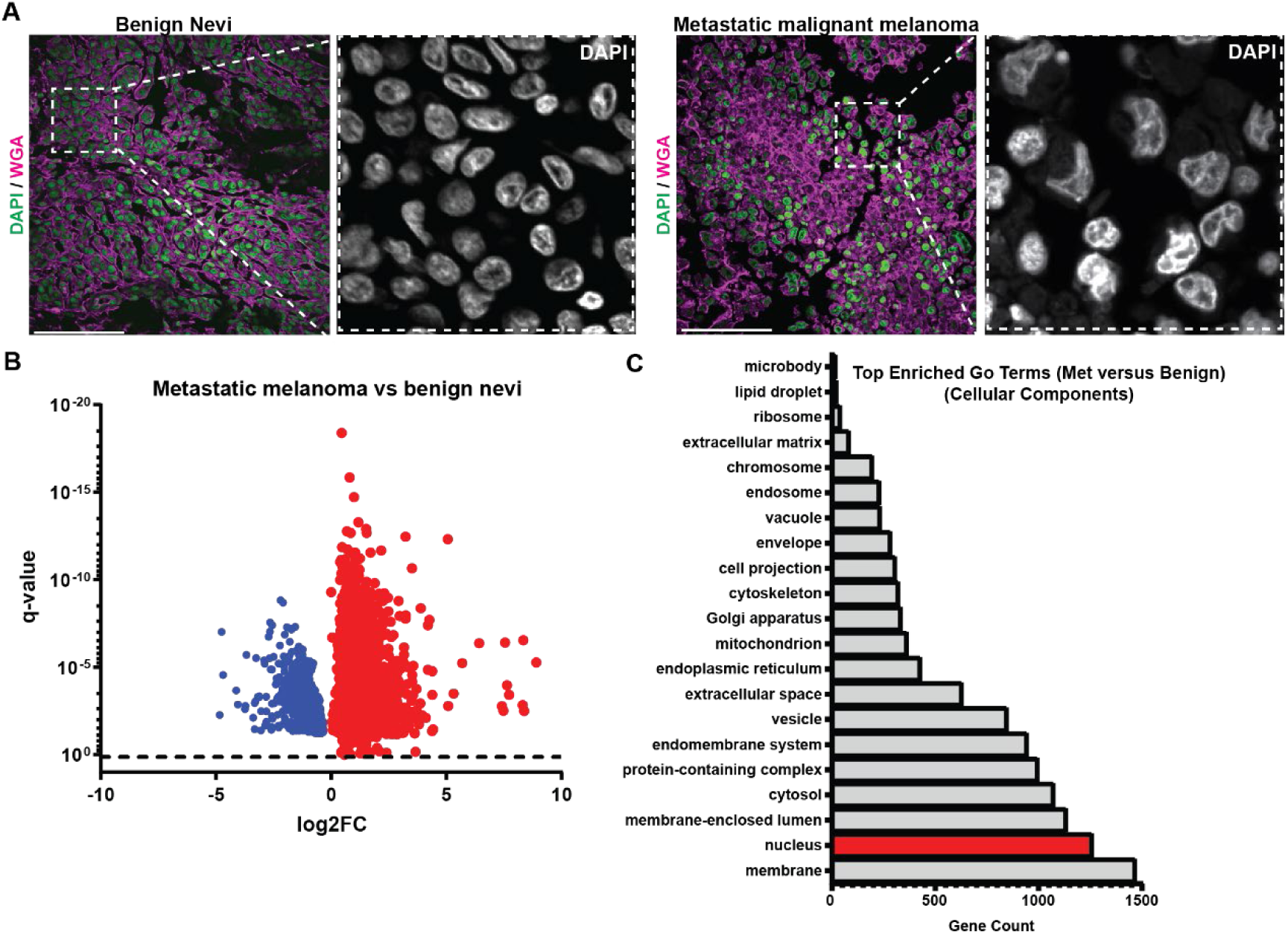
**(A)** Confocal images of tissue microarray sections of human clinical samples of benign nevi and metastatic melanoma, DNA stained with DAPI (left, green, and right grayscale) and the plasma membrane stained with Alexa 488 wheat germ agglutinin (WGA, left, purple). Bar= 100µm. **(B)** Volcano plot of *differential expression analysis* (red= significantly upregulated, blue= significantly downregulated) of genes in RNA-seq datasets (GEO:GSE98394) that differed in transcript abundance between patient biopsies from benign nevi and stage IV invasive tumors that were also present in genes differentially expressed between human melanocytes and 1205Lu cells, as well as from the transcriptomes of 27 melanoma cell lines from the Broad cancer cell line encyclopedia and from FACS-sorted patient melanoma cells (GEO:GSE72056). **(C)** Genes present in all four datasets were subjected to Gene ontology (GO) cellular component pathway analysis, red= nucleus associated genes.

**Fig. S3.**
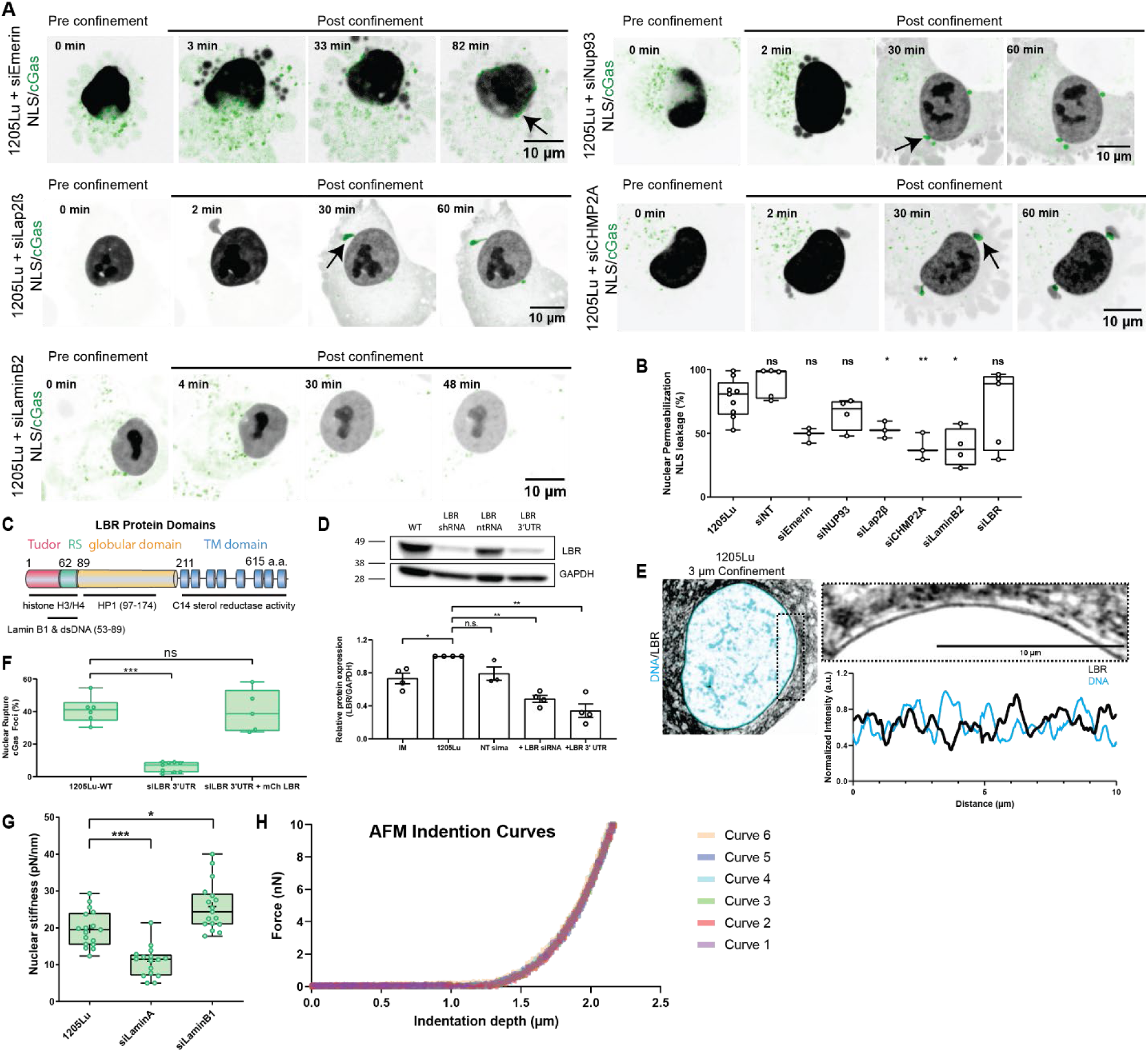
**(A)** Confocal image series of 1205Lu melanoma cells transfected with mCherry-NLS (inverted grayscale) and GFP-cGas (green) together with siRNAs targeting emerin (upper left), NUP93 (upper right), Lap2b (middle left), CHMP2A (middle right) or laminB2 (lower left), before and after confinement to 3µm. Arrows highlight cGas foci, bars= 10µm. **(B)** Quantification of the fraction of confined cells exhibiting NE permeabilization in 1205Lu cells with or without (1205Lu, n=9 experiments, 2473 cells) co-transfection with non-targeting siRNA (SiNT, n=5 experiments, 421 cells) or siRNA targeting emerin (siEmerin, n=3 experiments, 261 cells), NUP93 (siNUP93, n=4 experiments, 521 cells), Lap2β (siLap2β, n=3 experiments, 195 cells), CHMP2A (siCHMP2A, n=3 experiments, 204 cells), LaminB2 (siLaminB2, n=4 experiments, 607 cells), or LBR (siLBR, n=5 experiments, 1627 cells). **(C)** Schematic diagram of LBR functional domains, amino acid number noted. **(D)** Top: Western blot showing shRNA or shRNA 3’UTR mediated LBR depletion in 1205Lu cells. Bottom: Quantitative analysis of LBR levels from western blots of lysates of melanocytes (IM-WT) or 1205Lu cells with or without (1205Lu WT) transfection with non-targeting siRNAs (NT sirna), pooled siRNAs targeting the coding region of LBR (1205Lu LBR KD smartpool) or shRNAs targeting the 5’UTR of LBR (1205Lu LBR KD 3’UTR). **(E)** Super-resolution image taken during confinement to 3µm of a 1205Lu cell transfected with GFP-LBR and stained with SiR DNA (blue). Zoom of boxed region (above, right), intensity linescan along NE (below, right). **(F)** Quantification of the fraction of confined cells exhibiting NE rupture in 1205Lu cells with or without (1205Lu, n= 3 experiments, 152 cells) co-transfection with 3’UTR siRNA (siLBR 3’UTR, n= 3 experiments, n=3668 cells) or 3’UTR siRNA together with mCherry LBR (siLBR 3’UTR + mCherry LBR, n= 3 experiments, 1167 cells). **(G)** AFM quantification of nuclear stiffness in 1205Lu cells (n=7, n=43 cells), depleted for lamin A (n=3, n=16 cells), or lamin B1 (n=3, n=16 cells). **(H)** Example repeated force curve replicates performed on a single cell. In (B, F) points= mean of all cells from each imaging dish, in (D) points= individual experiments, in (G) points=individual cells. Significance was tested with a Student’s T-test Mann-Whitney (B,F), a paired students T-test (D), or a Student’s T-test Kolmogorov-Smirnov (G) bar= mean; error bars, SEM (E), bar= mean; error bars, min and max (B,F,G).

**Fig. S4.**
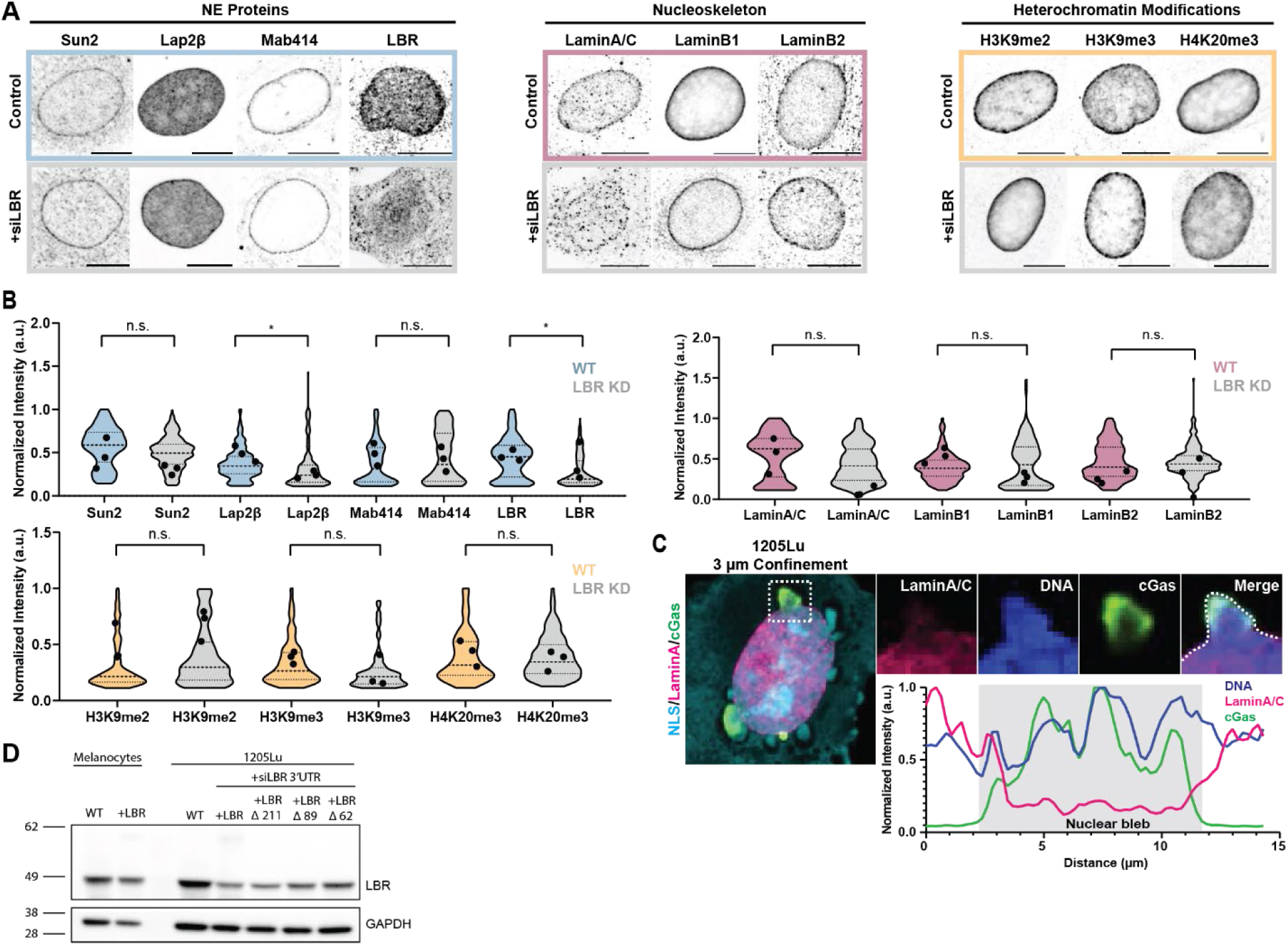
**(A)** Confocal images of 1205 Lu cells with (grey, +siLBR bottom rows) or without (top rows) transfection with siRNAs targeting LBR that were fixed and immunostained for NE proteins (left panel, blue, Sun2, Lap2β, nuclear pores (Mab414) and LBR), the nucleoskeleton (middle panel, dusky rose, lamin A/C, lamin B1, lamin B2) and heterochromatin modifications (right panel, melon, H3K9me2, H3K9me3, H4K20me3). **(B)** Quantification of immunostaining at the NE of images of cells treated as in (A). Sun2: WT n=3 experiments, 198 cells; siLBR n=3 experiments, 242 cells. Lap2β: WT n=3 experiments, 265 cells; siLBR n=3 experiments, 216 cells. Nuclear pores (Mab414): WT n=3 experiments, 242 cells; siLBR n=3 experiments, 116 cells. LBR: WT n=3 experiments, 268 cells; siLBR n=3 experiments, 173 cells. Lamin A/C: WT n=3 experiments, 112 cells; siLBR n=3 experiments, 175 cells. Lamin B1: WT n=3 experiments, 138 cells; siLBR n=3 experiments, 120 cells. Lamin B2: WT n=3 experiments, 273 cells; siLBR n=3 experiments, 220 cells. H3K9me2: WT n=3 experiments, 165 cells; siLBR n=3 experiments, 169 cells. H3K9me3: WT n=x3 experiments, 227 cells; siLBR n=3 experiments,108 cells. H4K20me3: WT n=3experiments, 138 cells; siLBR n=3 experiments, 114 cells. **(C)** Confocal image of a 1205Lu melanoma cell transfected with GFP-cGas (green) and mCherry NLS (cyan) that was fixed during confinement to 3µm and stained with SiR DNA (blue) and immunostained for Lamin A/C (magenta). Zoom of boxed region (above, right), intensity linescan along dotted line (below, right). **(D)** Western blot of LBR shRNA knockdown and rescue of LBR mutants. In (A) bar = 10µm. In (B), data points represent means of individual experiments. Significance was tested with a Student’s T-test Welch correction, bar= mean.

**Fig. S5.**
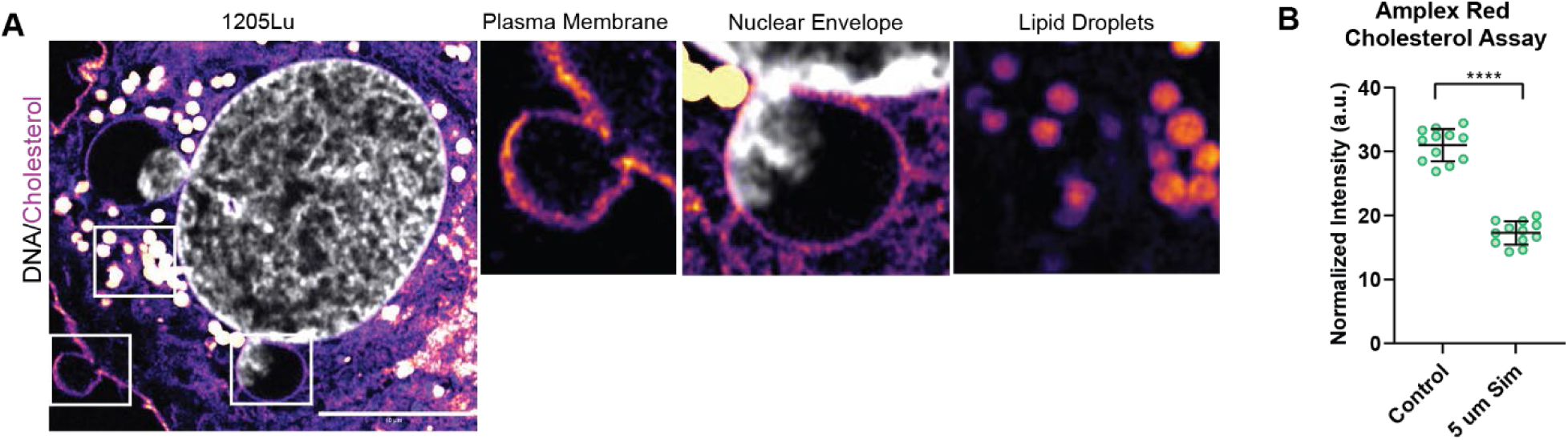
**(A)** Intensity color-encoded super-resolution confocal images taken during confinement to 3µm of 1205Lu melanoma cell stained with bodipy cholesterol and SiR DNA (grey). Zoom of boxed regions (above, right). **(B)** Spectrophotometric determination of cholesterol level in lysates of 1205Lu treated with DMSO (Vehicle) or with 5 µM simvastatin. N= 3 experiments, data point Mann-Whitney, bar= mean, error bars with SD. represent means of individual experiments. Significance was tested with a Student’s T-test Mann-Whitney, bar= mean, error bars with SD.

**Fig. S6.**
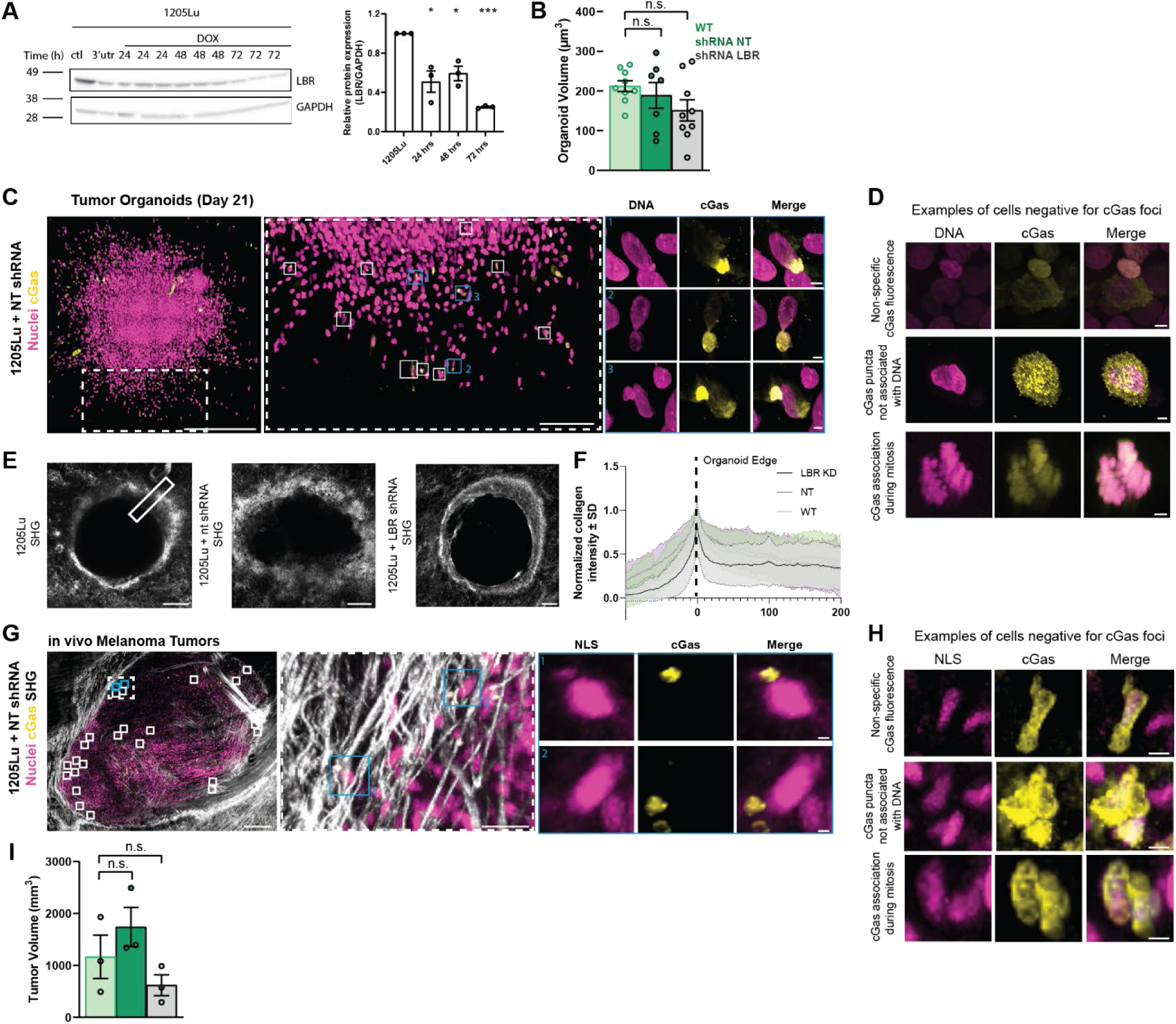
**(A)** Western blot of doxocycline induced stable knockdown of LBR. **(B)** Quantification of organoid volume) **(C)** 3D projections of z-series of confocal images of fixed tumor organoids grown from 1205Lu cells stably expressing GFP-cGAS (yellow) and mScarlet-NLS (magenta) with stable expression of non-targeting shRNA (+ntshRNA) for 21 days in a collagen ECM (bar = 500µm). Dotted boxed regions from left are zoomed (left center, bar = 150µm), white boxes indicate cells positive for cGas foci, blue boxed regions from left center are zoomed (right center, bar = 5µm). **(D)** Representative images of cells from organoids stably expressing GFP-cGAS (yellow) and mScarlet-NLS (magenta) quantified as negative for cGas foci. (bar = 5µm). **(E)** 3D projections of z-series of two-photon images of organoids grown from 1205Lu cells with either (shRNA LBR, n=3 organoids) or (NT shRNA, n=3 organoids) or without (1205Lu, n=3 organoids). Box indicates line scan region and thickness. **(F)** Quantification of collagen intensity from (E). **(G)** 3D projection of z-series of two-photon images of *in vivo* tumors grown from 1205Lu cells stably expressing GFP-cGAS (yellow) and mScarlet-NLS (magenta) with (NT shRNA) in mouse dermis (bar=500µm). Solid boxed regions from left represent nuclei positive for cGas foci and blue boxes are zoomed (right, bar=50µm), dashed boxed region from left is zoomed (center, bar=50µm). **(H)** Representative images of cells from tumors stably expressing GFP-cGAS (yellow) and mScarlet-NLS (magenta) quantified as negative for cGas foci (bar = 5µm). Significance was tested with a paired student’s T-test (A); bars=mean, error bars with SEM. In (B) significance was tested with a Student’s T-test Welch correction, bars= means with SEM.

**Fig. S7.**
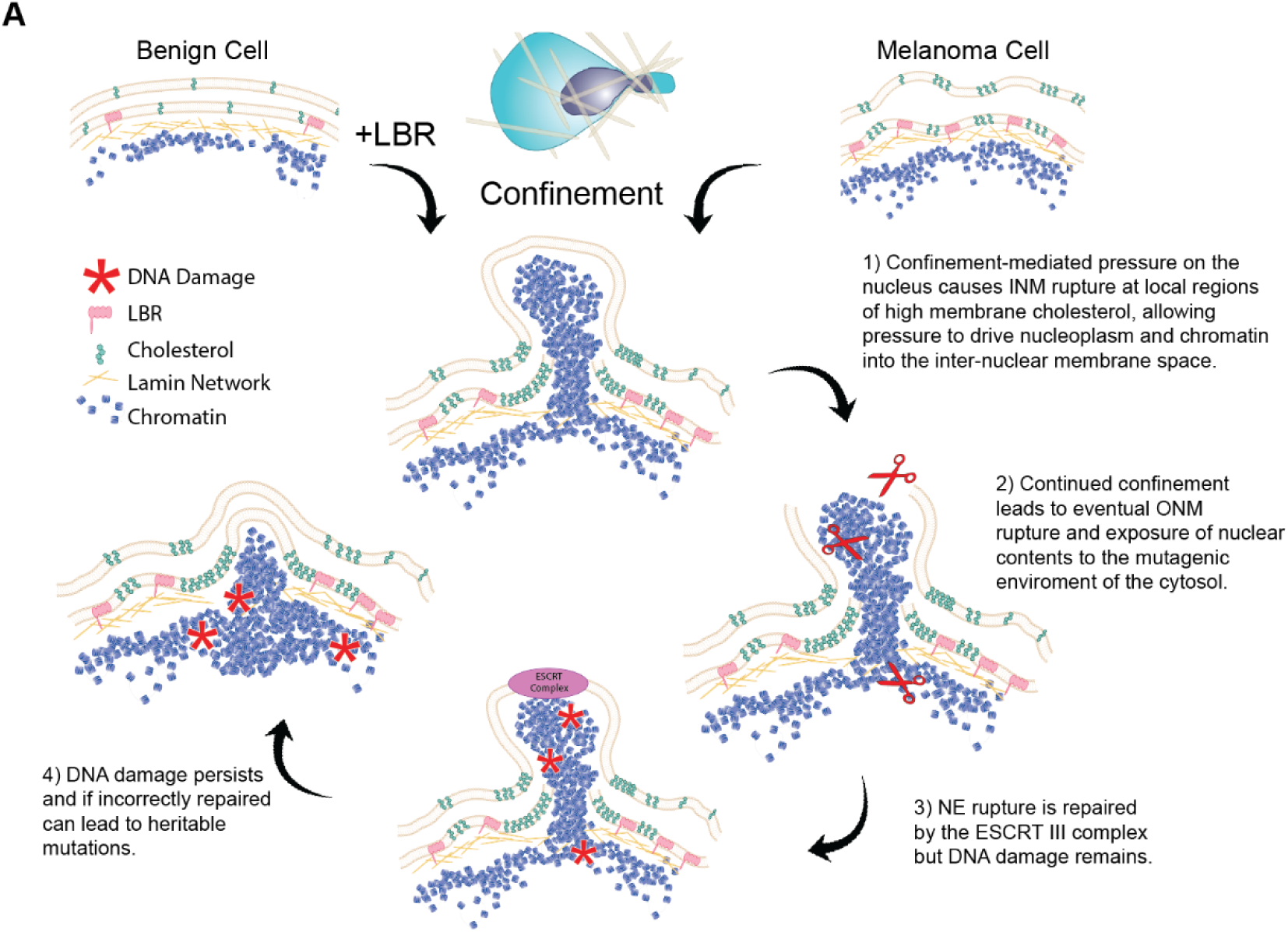
**(A)** Model summarizing how LBR-mediated cholesterol accumulation in the NE promotes NE rupture during confined migration.

**Movie S1.**

Confocal image series of benign and malignant cells before and after confinement to 3µm. Representative cell lines for skin (immortalized human melanocytes, 1205Lu), breast (MCF10A or T47D) and prostate (RWPE-1, PC-3) transfected with mCherry-NLS (inverted grayscale) and GFP-cGas (green). Arrows highlight GFP-cGas foci indicating nuclear rupture. Time interval between images 60 s. Scale bar, 10µm.

**Movie S2.**

Confocal image series of 1205Lu cells transfected with mCherry-NLS (inverted grayscale) and GFP-cGas (green) with and without treatment with the following siRNA’s to specifically deplete NE proteins; 1205Lu WT (top, left), nt siRNA (top, center left), emerin (top, center right), NUP93 (top right), Lap2b (bottom left), CHMP2A (bottom center left), laminB2 (bottom center right) or LBR (bottom right), before and after confinement to 3mm. Arrows highlight GFP-cGas foci indicating nuclear rupture. Time interval between images 120 s. Scale bars, 10µm.

**Movie S3.**

Confocal image series of immortalized human melanocytes transfected with mCherry-NLS or mCherry-LBR (inverted grayscale) and GFP-cGas (green) before and after confinement to 3µm. Arrows highlight GFP-cGas foci indicating nuclear rupture. Time interval between images 120 s. Scale bar, 10µm.

**Movie S4.**

Superresolution Z-series of a 1205Lu transfected with EGFP-Lap2b (magenta), and vital dyes blue-white ER tracker (green) and siR-DNA (blue) after confinement to 3mm, highlighting differential rupture of the INM and ONM. Scale bar as shown in movie. Superresolution Z-series of a 1205Lu transfected with siRNA depleting LBR, EGFP-Lap2b (magenta), and vital dyes blue-white ER tracker (green) and siR-DNA (blue) after confinement to 3mm, highlighting differential rupture of the INM and ONM. Scale bar as shown in movie.

**Movie S5.**

Confocal image series of 1205Lu cells transfected with GFP-cGas (green) and treated and siRNA targeting the 3’UTR of LBR (+LBR siRNA) together with either mCherry-LBR (left, inverted grayscale), mCherry-LBRDTudor+RS (left center, inverted grayscale), mCherry LBR 583Q (right center, inverted grayscale) or mCherry LBR 1402DT (right end, inverted grayscale), before and after confinement to 3mm. Arrows highlight GFP-cGas foci indicating nuclear rupture. Time interval between images 120 s. Scale bar, 10µm.

**Movie S6.**

Confocal image series of 1205Lu cells transfected with mCherry-NLS (inverted grayscale) and GFP-cGas (green) treated with either vehicle control (DMSO) or 20mM simvastatin, before and after confinement to 3mm. Arrow indicates cGas focus, indicating a cell positive for nuclear rupture. Time interval between images 120 s. Scale bar, 10µm.

**Movie S7.**

3D reconstruction of a 17.5µM-thick confocal Z-series of fixed tumor organoids grown from 1205Lu cells stably expressing GFP-cGas (yellow) and mScarlet-NLS (magenta) in a collagen ECM. Scale bar as shown in movie.

**Movie S8.**

3D reconstruction of a 36µM-thick confocal Z-series of fixed tumor organoids grown from 1205Lu cells stably expressing GFP-cGas (yellow), mScarlet-NLS (magenta) and shRNA targeting LBR, in a collagen ECM. Scale bar as shown in movie.

**Movie S9.**

3D reconstruction of a 223µM-thick two-photon confocal Z-series of living ex-vivo tumors grown from 1205Lu cells stably expressing GFP-cGAS (yellow) and mScarlet-NLS (magenta) in mouse dermis. Scale bar as shown in movie. Gamma correction was applied to SHG channel (greyscale) to allow for better visualization of dim collagen fibers.

**Movie S10.**

3D reconstruction of a 145µM-thick two-photon confocal Z-series of living ex-vivo tumors grown from 1205Lu cells stably expressing GFP-cGAS (yellow), mScarlet-NLS (magenta) and shRNA targeting LBR in mouse dermis. Scale bar as shown in movie. Gamma correction was applied to SHG channel (greyscale) to allow for better visualization of dim collagen fibers.

