## Supplemental Files for "Lamin B Receptor Upregulation in Metastatic Melanoma Causes Cholesterol-Mediated Nuclear Envelope Fragility"

Supplement 1

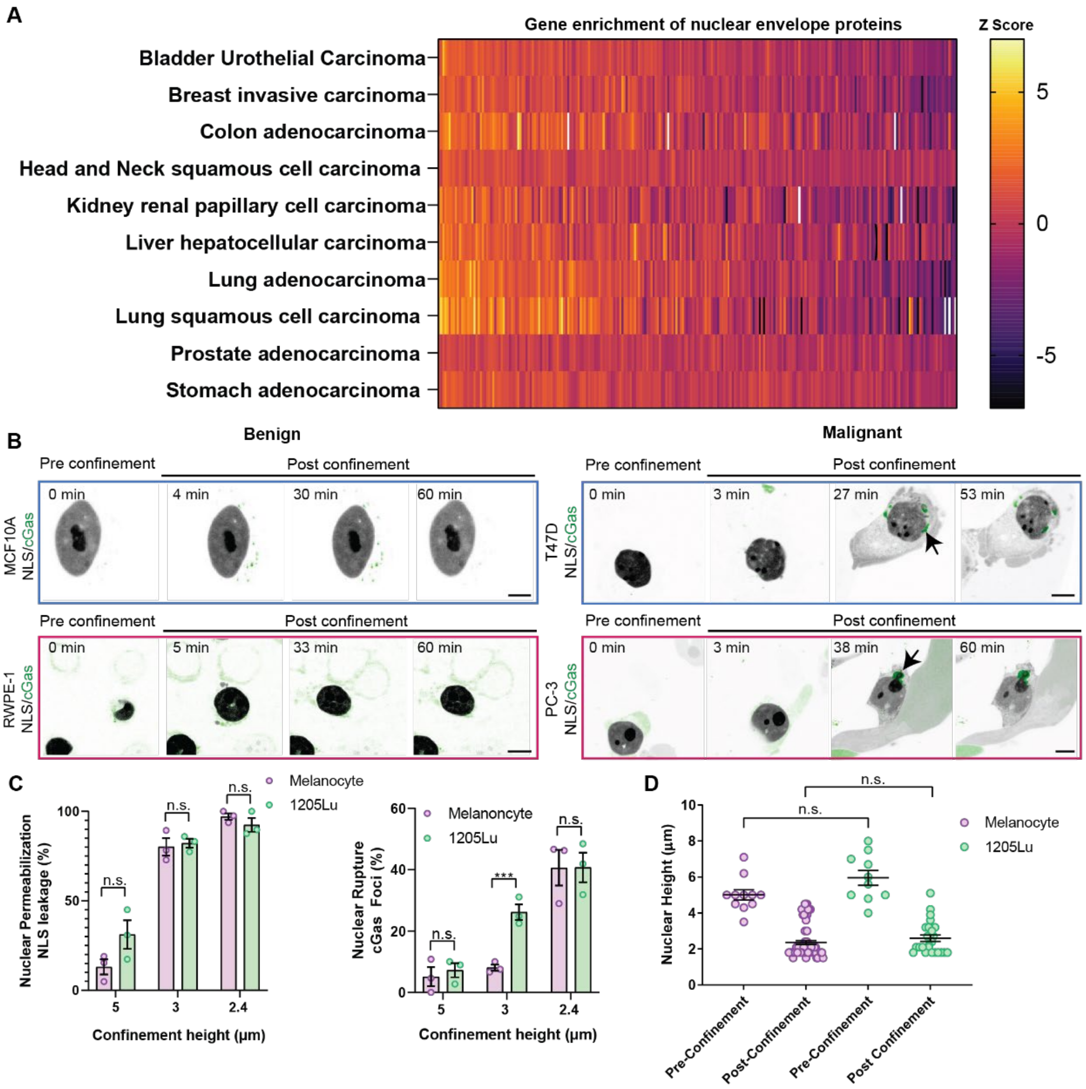


### Fig. S1.

**(A)** Heat map of transcript abundance (normalized relative to a baseline human genome) of 249 genes encoding nuclear envelope proteins (curated from the Human Protein Atlas) from RNA-seq data from 6860 clinical samples representing 10 cancer subtypes from The Cancer Genome Atlas (TCGA) PanCancer repository. **(B)** Confocal image series of living MCF10A (upper left) or T47D (upper right) breast cells (blue outline) or RWPE-1 (lower left) or PC-3 (lower right) prostate cells (pink outline) transfected with mCherry-NLS (inverted grayscale) and GFP-cGas (green) before and after confinement to 3μm. Arrows highlight cGas foci. Bar = 10μm. **(C)** Quantification of NE permeabilization (left) and NE rupture (right) in human melanocytes (purple) and 1205Lu cells (green) at different confinement heights (Melanocytes n=3 experiments for all heights, n=404 cells (2.4μm), n=196 cells (3μm), n=184 cells (5μm); 1205Lu n=3 experiments for all heights, n=301 cells (2.4μm) n=604 cells (3μm), n=1375 cells (5μm). **(D)** Nuclear height of fixed 1205Lu cells (green) and human melanocytes (purple) expressing GFP-NLS measured from confocal image series before and after confinement to 3μm. 1205Lu: before confinement n=3 experiments, 189 cells; after confinement n=3 experiments, 159 cells. Human melanocytes: before confinement n=3 experiments, 107 cells;

after confinement n=3 experiments, 71 cells. Data points represent independent experiments each consisting of pooled cell data from a single imaging dish (C) or individual cells (D). Significance was tested with a Student's T-test Mann-Whitney, bar= mean, error bars= SEM.

Supplement 2

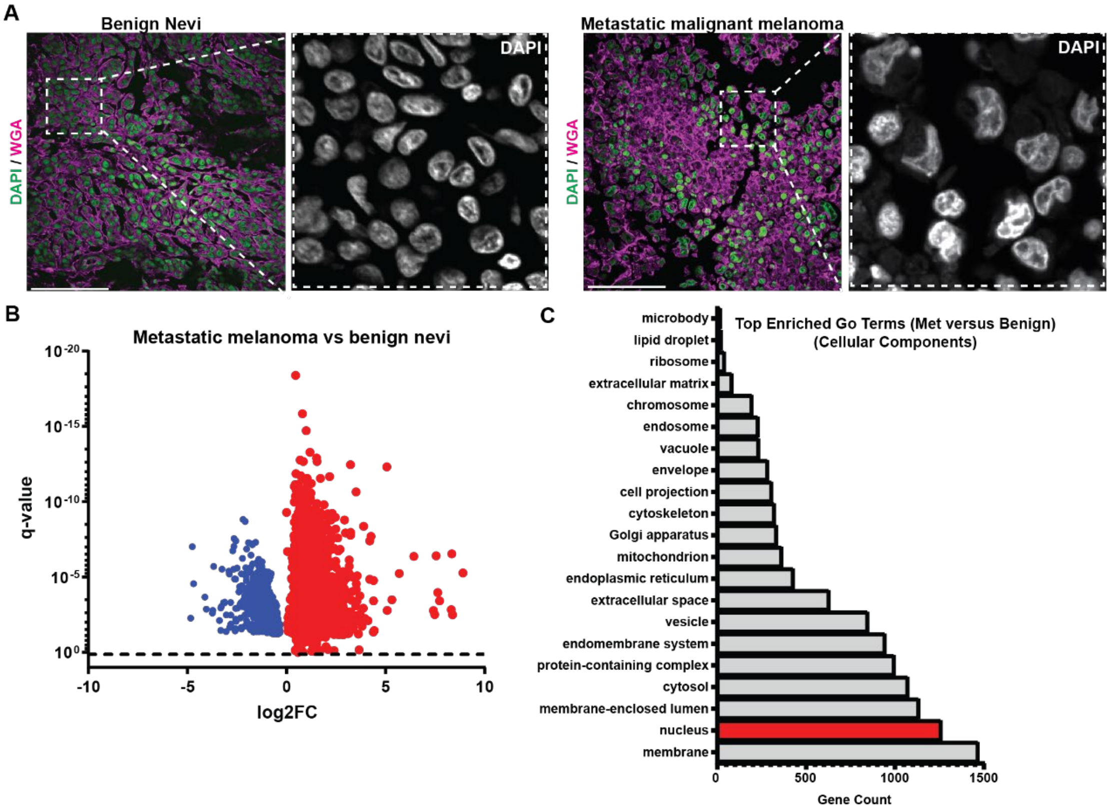


**Fig. S2.**

**(A)** Confocal images of tissue microarray sections of human clinical samples of benign nevi and metastatic melanoma, DNA stained with DAPI (left, green, and right grayscale) and the plasma membrane stained with Alexa 488 wheat germ agglutinin (WGA, left, purple). Bar= 100μm. **(B)** Volcano plot of *differential expression analysis* (red= significantly upregulated, blue= significantly downregulated) of genes in RNA-seq datasets (GEO:GSE98394) that differed in transcript abundance between patient biopsies from benign nevi and stage IV invasive tumors that were also present in genes differentially expressed between human melanocytes and 1205Lu cells, as well as from the transcriptomes of 27 melanoma cell lines from the Broad cancer cell line encyclopedia and from FACS-sorted patient melanoma cells (GEO:GSE72056). **(C)** Genes present in all four datasets were subjected to Gene ontology (GO) cellular component pathway analysis, red= nucleus associated genes.

Supplemental Figure 3

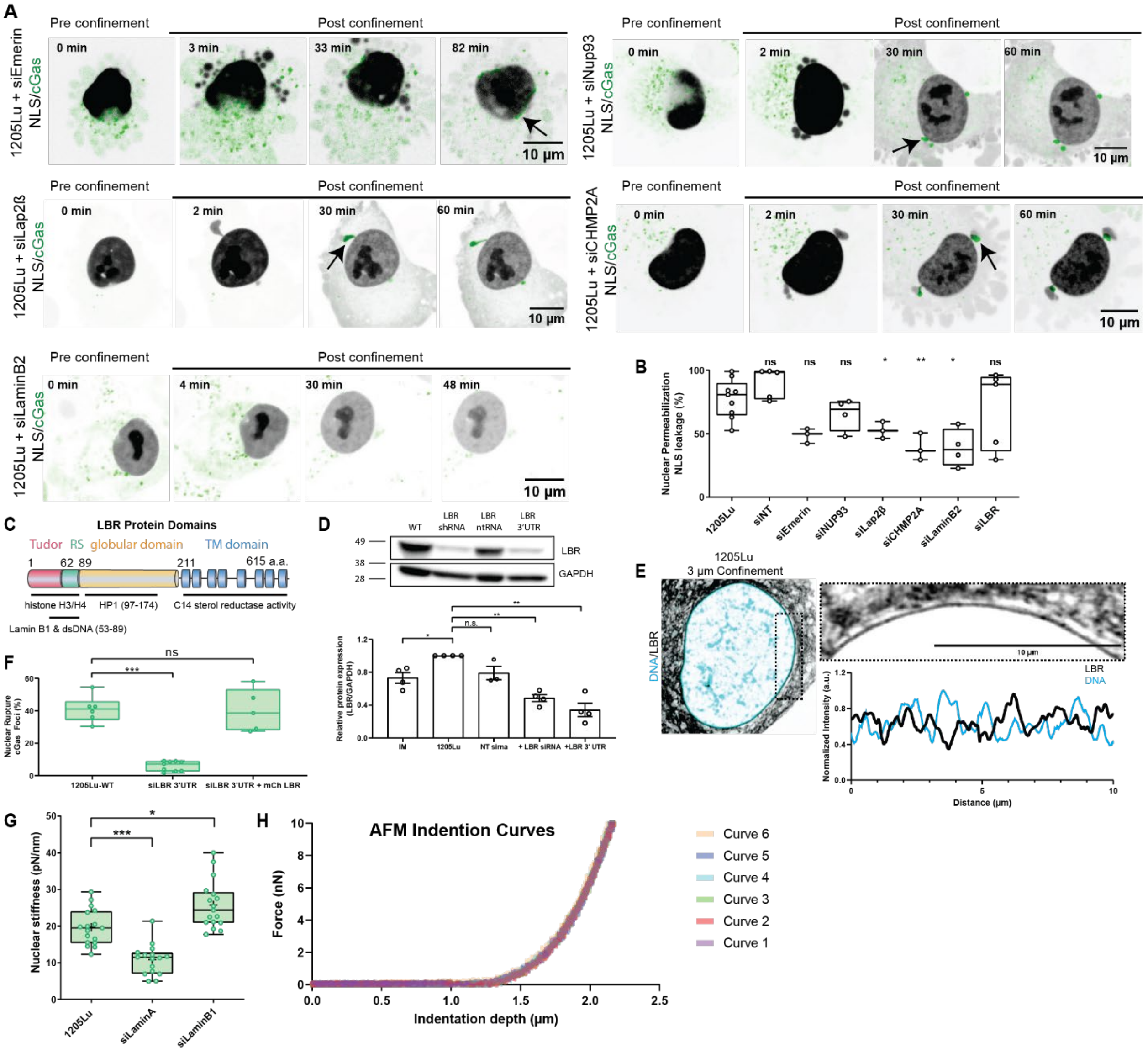

**Fig. S3.**

**(A)** Confocal image series of 1205Lu melanoma cells transfected with mCherry-NLS (inverted grayscale) and GFP-cGas (green) together with siRNAs targeting emerin (upper left), NUP93 (upper right), Lap2b (middle left), CHMP2A (middle right) or laminB2 (lower left), before and after confinement to 3µm. Arrows highlight cGas foci, bars= 10µm. **(B)** Quantification of the fraction of confined cells exhibiting NE permeabilization in 1205Lu cells with or without (1205Lu, n=9 experiments, 2473 cells) co-transfection with non-targeting siRNA (SiNT, n=5 experiments, 421 cells) or siRNA targeting emerin (siEmerin, n=3 experiments, 261 cells), NUP93 (siNUP93, n=4 experiments, 521 cells), Lap2β (siLap2β, n=3 experiments, 195 cells), CHMP2A (siCHMP2A, n=3 experiments, 204 cells), LaminB2 (siLaminB2, n=4 experiments, 607 cells), or LBR (siLBR, n=5 experiments, 1627 cells). **(C)** Schematic diagram of LBR functional domains, amino acid number noted. **(D)** Top: Western blot showing shRNA or shRNA 3'UTR mediated LBR depletion in 1205Lu cells. Bottom: Quantitative analysis of LBR levels from western blots of lysates of melanocytes (IM-WT) or 1205Lu cells with or without (1205Lu WT) transfection with non-targeting siRNAs (NT sirna), pooled siRNAs targeting the coding region of LBR (1205Lu

LBR KD smartpool) or shRNAs targeting the 5'UTR of LBR (1205Lu LBR KD 3'UTR). **(E)** Super-resolution image taken during confinement to 3µm of a 1205Lu cell transfected with GFP-LBR and stained with SiR DNA (blue). Zoom of boxed region (above, right), intensity linescan along NE (below, right). **(F)** Quantification of the fraction of confined cells exhibiting NE rupture in 1205Lu cells with or without (1205Lu, n= 3 experiments, 152 cells) co-transfection with 3'UTR siRNA (siLBR 3'UTR, n= 3 experiments, n=3668 cells) or 3'UTR siRNA together with mCherry LBR (siLBR 3'UTR + mCherry LBR, n= 3 experiments, 1167 cells). **(G)** AFM quantification of nuclear stiffness in 1205Lu cells (n=7, n=43 cells), depleted for lamin A (n=3, n=16 cells), or lamin B1 (n=3, n=16 cells). **(H)** Example repeated force curve replicates performed on a single cell. In (B, F) points= mean of all cells from each imaging dish, in (D) points= individual experiments, in (G) points=individual cells. Significance was tested with a Student's T-test Mann-Whitney (B,F), a paired students T-test (D), or a Student's T-test Kolmogorov-Smirnov (G) bar= mean; error bars, SEM (E), bar= mean; error bars, min and max (B,F,G).

Supplemental Figure 4

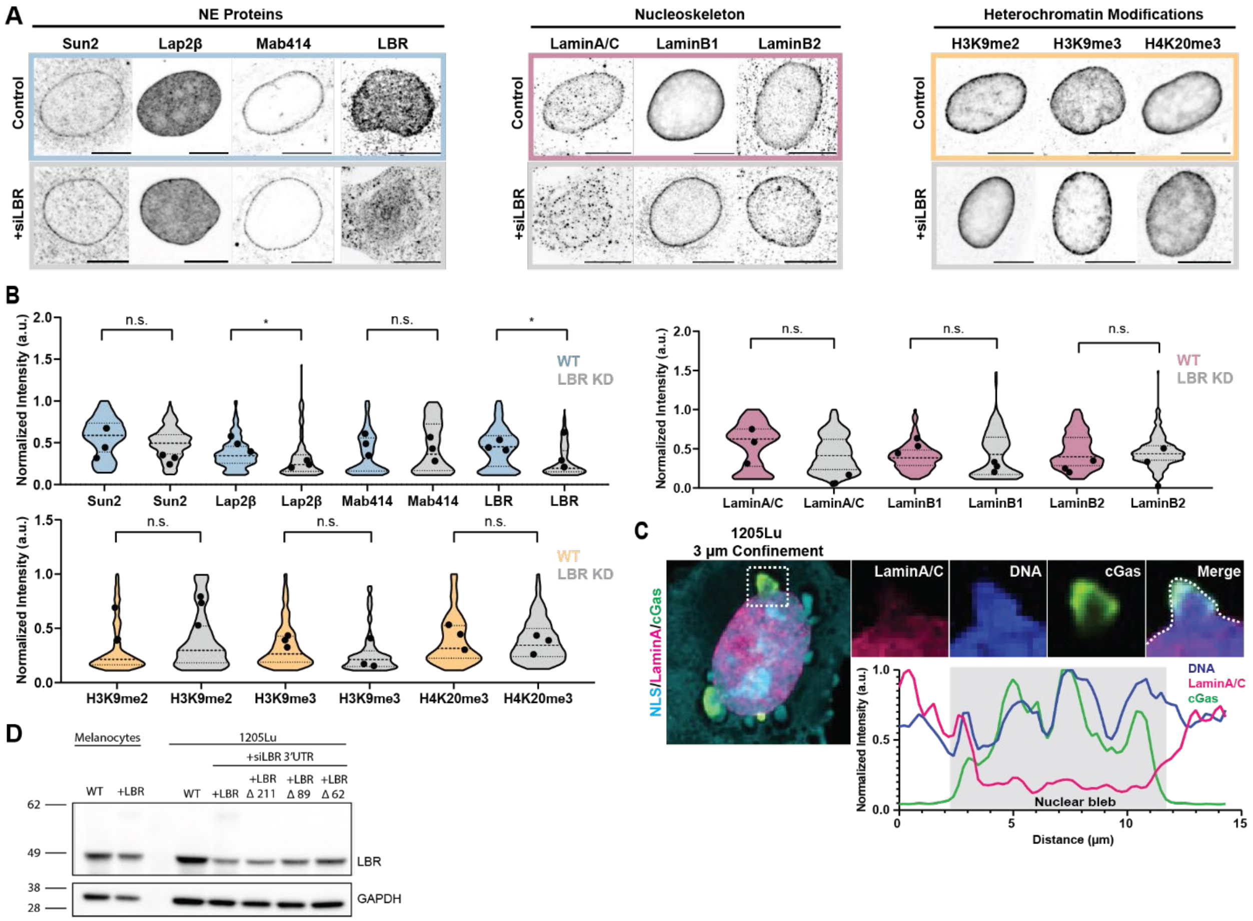


**Fig. S4. (A)** Confocal images of 1205 Lu cells with (grey, +siLBR bottom rows) or without (top rows) transfection with siRNAs targeting LBR that were fixed and immunostained for NE proteins (left panel, blue, Sun2, Lap2β, nuclear pores (Mab414) and LBR), the nucleoskeleton (middle panel, dusky rose, lamin A/C, lamin B1, lamin B2) and heterochromatin modifications (right panel, melon, H3K9me2, H3K9me3, H4K20me3). **(B)** Quantification of immunostaining at the NE of images of cells treated as in (A). Sun2: WT n=3 experiments, 198 cells; siLBR n=3 experiments, 242 cells. Lap2β: WT n=3 experiments, 265 cells; siLBR n=3 experiments, 216 cells. Nuclear pores (Mab414): WT n=3 experiments, 242 cells; siLBR n=3 experiments, 116 cells. LBR: WT n=3 experiments, 268 cells; siLBR n=3 experiments, 173 cells. Lamin A/C: WT n=3 experiments, 112 cells; siLBR n=3 experiments, 175 cells. Lamin B1: WT n=3 experiments, 138 cells; siLBR n=3 experiments, 120 cells. Lamin B2: WT n=3 experiments, 273 cells; siLBR n=3 experiments, 220 cells. H3K9me2: WT n=3 experiments, 165 cells; siLBR n=3 experiments, 169 cells. H3K9me3: WT n=x3 experiments, 227 cells; siLBR n=3 experiments,108 cells. H4K20me3: WT n=3experiments, 138 cells; siLBR n=3 experiments, 114 cells. **(C)** Confocal image of a 1205Lu melanoma cell transfected with GFP-cGas (green) and mCherry NLS (cyan) that was fixed during confinement to 3µm and stained with SiR DNA (blue) and immunostained for Lamin A/C (magenta). Zoom of boxed region (above, right), intensity linescan along dotted line (below, right). **(D)** Western blot of LBR shRNA knockdown and rescue of LBR mutants. In (A) bar = 10µm. In (B), data points represent means of individual experiments. Significance was tested with a Student's T-test Welch correction, bar= mean.

Supplemental Figure 5

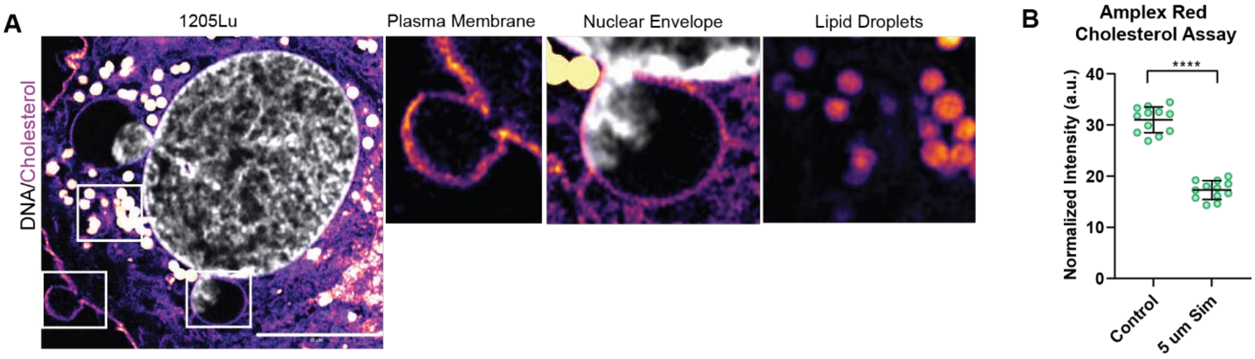


**Fig. S5.(A)** Intensity color-encoded super-resolution confocal images taken during confinement to 3μm of 1205Lu melanoma cell stained with bodipy cholesterol and SiR DNA (grey). Zoom of boxed regions (above, right). **(B)** Spectrophotometric determination of cholesterol level in lysates of 1205Lu treated with DMSO (Vehicle) or with 5 μM simvastatin. N= 3 experiments, data point Mann-Whitney, bar= mean, error bars with SD. represent means of individual experiments. Significance was tested with a Student's T-test Mann-Whitney, bar= mean, error bars with SD.

Supplemental Figure 6

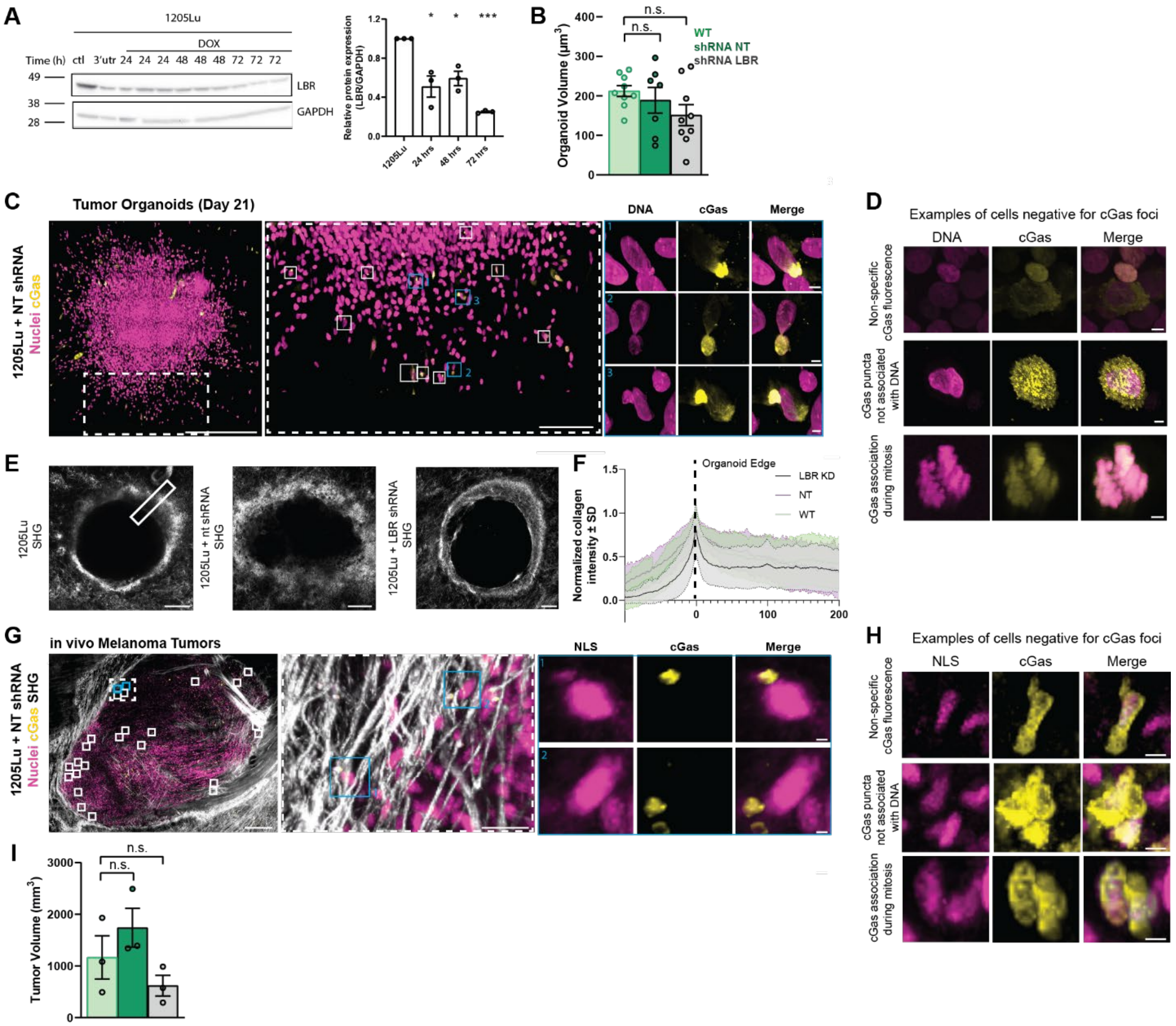


**Fig. S6**. **(A)** Western blot of doxocycline induced stable knockdown of LBR. **(B)** Quantification of organoid volume) **(C)** 3D projections of z-series of confocal images of fixed tumor organoids grown from 1205Lu cells stably expressing GFP-cGAS (yellow) and mScarlet-NLS (magenta) with stable expression of non-targeting shRNA (+ntshRNA) for 21 days in a collagen ECM (bar = 500μm). Dotted boxed regions from left are zoomed (left center, bar = 150μm), white boxes indicate cells positive for cGas foci, blue boxed regions from left center are zoomed (right center, bar = 5μm). **(D)** Representative images of cells from organoids stably expressing GFP-cGAS (yellow) and mScarlet-NLS (magenta) quantified as negative for cGas foci. (bar = 5μm). **(E)** 3D projections of z-series of two-photon images of organoids grown from 1205Lu cells with either (shRNA LBR, n=3 organoids) or (NT shRNA, n=3 organoids) or without (1205Lu, n=3 organoids). Box indicates line scan region and thickness. **(F)** Quantification of collagen intensity from (E). **(G)** 3D projection of z-series of two-photon images of *in vivo* tumors grown from 1205Lu cells stably expressing GFP-cGAS (yellow) and mScarlet-NLS (magenta) with (NT shRNA) in mouse dermis (bar=500μm). Solid boxed regions from left represent nuclei positive for cGas foci and blue boxes are zoomed (right, bar=50μm), dashed boxed region from left is zoomed (center, bar=50µm). **(H)** Representative images of cells from tumors stably expressing GFP-cGAS (yellow) and mScarlet-NLS (magenta) quantified as negative for cGas foci (bar = 5µm). Significance was tested with a paired student's T-test (A); bars=mean, error bars with SEM. In (B) significance was tested with a Student's T-test Welch correction, bars= means with SEM.

Supplemental Figure 7

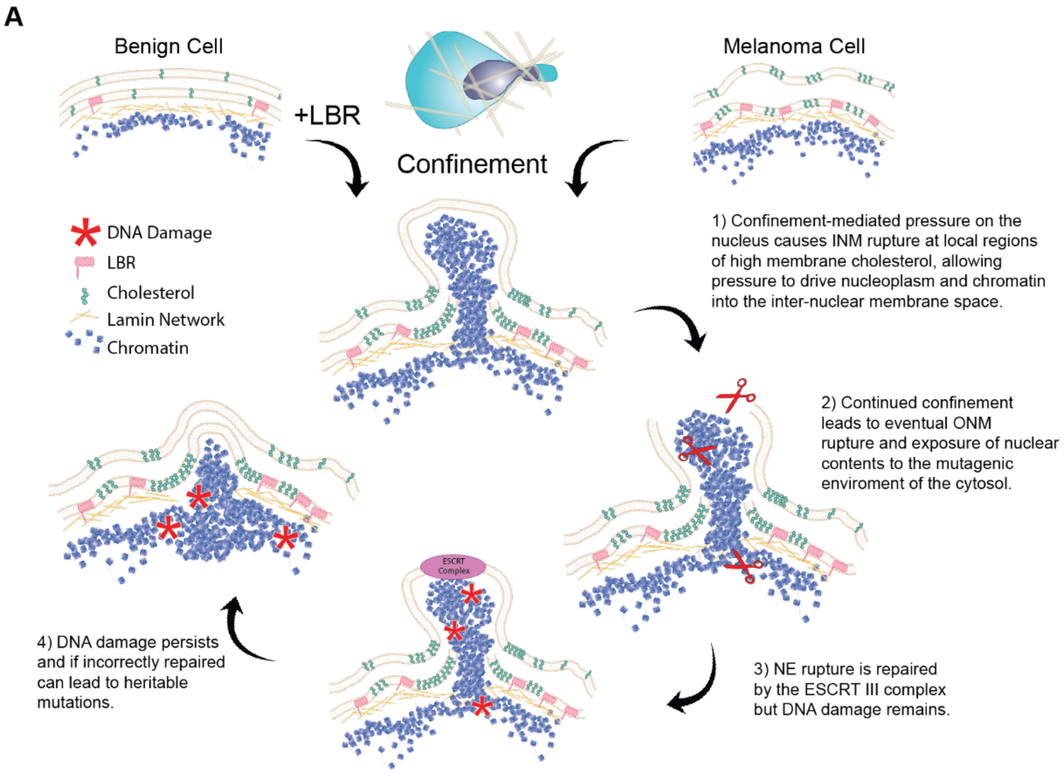


**Fig. S7**.
**(A)** Model summarizing how LBR-mediated cholesterol accumulation in the NE promotes NE rupture during confined migration.

**Movie S1.**
Confocal image series of benign and malignant cells before and after confinement to 3µm. Representative cell lines for skin (immortalized human melanocytes, 1205Lu), breast (MCF10A or T47D) and prostate (RWPE-1, PC-3) transfected with mCherry-NLS (inverted grayscale) and GFP-cGas (green). Arrows highlight GFP-cGas foci indicating nuclear rupture. Time interval between images 60 s. Scale bar, 10µm.
